# Embryoid bodies facilitate comparative analysis of gene expression in humans and chimpanzees across dozens of cell types

**DOI:** 10.1101/2022.07.20.500831

**Authors:** Kenneth A Barr, Katherine L Rhodes, Yoav Gilad

## Abstract

Comparative gene expression studies in apes are fundamentally limited by the challenges associated with sampling across different tissues. Here, we used single-cell RNA-sequencing of embryoid bodies (**EBs**) to collect transcriptomic data from over 70 cell types in three humans and three chimpanzees. We found hundreds of genes whose regulation is conserved across cell types, as well as genes whose regulation likely evolves under directional selection in one or a handful of cell types. Using EBs from a human-chimpanzee fused cell line, we also inferred the proportion of inter-species regulatory differences due to changes in *cis* and *trans* elements between the species. Thus, we present the most comprehensive dataset of comparative gene expression from humans and chimpanzees to date, including a catalog of regulatory mechanisms associated with inter-species differences.

## Introduction

Comparative functional genomic studies in primates reveal insight into the evolution of gene regulation and help us identify genes and pathways that are associated with species-specific traits (Anderson et al., 2020; Berthelot et al., 2018; Blake et al., 2020; Eres et al., 2019; Fair et al., 2020; Housman & Gilad, 2020; Klein et al., 2018; Mittleman et al., 2021). In particular, studies of gene expression in humans and other apes allow us to identify regulatory changes that may be associated with human-specific adaptations (Danko et al., 2018; Housman & Gilad, 2020; O’Bleness et al., 2012; Pizzollo et al., 2018). Comparative inter-species studies also allow us to identify patterns of regulatory variation that are consistent with the action of natural selection. Such patterns suggest functional importance and point to regulatory phenotypes that may affect fitness. The inference of fitness-related molecular function is profoundly important, not only to studies that attempt to understand the mechanisms of evolutionary change, but also to studies of the genetic and gene regulatory basis for complex traits and diseases in humans (Benito-Kwiecinski et al., 2021; Fair et al., 2020; Gokhman et al., 2021; Housman et al., 2022; Mittleman et al., 2021; Pizzollo et al., 2018; Pollen et al., 2019; Shibata et al., 2012; Ward & Gilad, 2019).

One goal of comparative genomic studies in primates has been to characterize gene regulatory variation across a wide range of tissue types. Comparative genomic studies are not typically hypothesis-driven; rather, they aim to build comprehensive comparative catalogs that can be used to infer function, explore the evolution of different regulatory mechanisms, and establish hypotheses based on causal inference between genotypes and phenotypes (Berthelot et al., 2018; Blake et al., 2020; Danko et al., 2018; Eres et al., 2019; Fair et al., 2020; Klein et al., 2018; Mittleman et al., 2021; Pizzollo et al., 2018). Yet, more than two decades after the first genome-scale comparison of gene expression in humans and chimpanzees (Enard et al., 2002), comparative genomic data from apes remains limited to just a handful of tissue types (Housman & Gilad, 2020).

For obvious ethical and practical reasons, only a small number of tissue types can be accessed from live humans, and direct experimentation *in vivo* is impossible. While *in vivo* studies are permitted in some primates, ethical considerations forcefully apply to studies of apes, particularly to studies involving chimpanzees – our closest extant evolutionary relatives. Comparative studies in apes must therefore rely on opportunistic collection of post-mortem tissues. Efforts to collect a broad array of post-mortem tissues from humans have been quite successful (GTEx Consortium, 2020), but comparative genomic studies in humans and chimpanzees have only examined about a dozen different tissues. Moreover, the sample size in nearly all comparative studies in apes is quite modest: most studies include just 4-6 donors from each non-human species (Barreiro et al., 2010; Blake et al., 2018, 2020; Blekhman et al., 2008, 2010; Cáceres et al., 2003; Cain et al., 2011; De la Cruz et al., 2009; Eres et al., 2019; Gilad et al., 2006; Hernando-Herraez et al., 2013; Housman et al., 2022; Iskow et al., 2012; Khan et al., 2013; Lemos et al., 2005; Mittleman et al., 2021; Pai et al., 2011; Perry et al., 2012), and in some cases, only a single sample from the non-human species was available (Brawand et al., 2011; Enard et al., 2002; Gilad et al., 2005; Hernando-Herraez et al., 2015). In the handful of comparative studies that examine more than one tissue in apes, multiple tissues are rarely sampled from the same individuals (Barbosa-Morais et al., 2012; Blekhman et al., 2008; Brawand et al., 2011; Cáceres et al., 2003; De la Cruz et al., 2009; Enard et al., 2002; Pai et al., 2011; Perry et al., 2012), resulting in a severe confounding of tissue and individual effects on the observed variation in gene regulation (a confounding effect that is particularly severe due to the small sample size available from each tissue (Blake et al., 2020)).

Even if we had unlimited access to post-mortem frozen tissues from apes, we would still be unable to study dynamic gene regulation, let alone study how different tissues and cell types respond to environmental exposures. Induced pluripotent stem cells (**iPSCs**) from apes provide a way to study gene regulatory dynamics during differentiation or in response to external perturbations (Gallego Romero et al., 2015), but the inefficiency of differentiation protocols limits the number of individuals, cell types, and contexts that can be queried in a single experiment. Thus, the scope of comparative studies in apes has remained narrow: we are unable to study gene regulation in more than a few tissues from apes; we are generally unable to collect samples from enough individuals to map and study regulatory quantitative trait loci (**QTLs**) in non-human apes; we are unable to study the dynamics of gene regulation during development; and we are unable to study regulatory responses to evolutionarily and clinically relevant exposures.

If we are ever to unleash the power of the comparative paradigm in apes, we require an *in vitro* system that allows us to dynamically study gene regulation in dozens of different cell types and individuals, and in hundreds of different contexts. Here, we show that embryoid bodies (**EBs**) can potentially fulfill these requirements. EBs are dynamic, iPSC-derived organoids that contain spontaneously and asynchronously differentiating cells from all three germ layers (Brickman & Serup, 2017).

We recently used single-cell RNA-sequencing (**scRNA-seq**) to characterize cellular and gene expression heterogeneity in a panel of human EBs, which revealed the presence of dozens of differentiating and terminal cell types from multiple different tissues (Rhodes et al., 2022). In the present study, we applied scRNA-seq to EBs from three humans and three chimpanzees. Despite the small size of our sample, we were able to assemble a comparative catalog of gene expression data from more than 70 different cell types per species, including differentiating cell types that have never been accessible from frozen tissues. This catalog contains nearly an order of magnitude more cell types than have been examined before, and three times more comparative data from humans and chimpanzees than all previous studies combined.

Using gene expression data from EBs, we identify hundreds of genes that are differentially expressed across dozens of cell types, as well as genes that are broadly conserved. We analyze tissue-restricted patterns of differential expression between the species, implicating a handful of candidate genes that may underlie functional differences between humans and chimpanzees. In addition, we quantify the contribution of *cis* and *trans* changes to inter-species differences in gene expression using EBs derived from a fused human-chimpanzee iPSC line.

## Results

### Generation of human and chimpanzee embryoid bodies

We generated EBs from three human YRI (Yoruba in Ibadan, Nigeria) and three western chimpanzee iPSC lines that were previously validated (Banovich et al., 2018; Gallego Romero et al., 2015). From each line, we established EBs in three independent replicates (**Figure 1A and Figure S1**; see Materials and methods). We also included a single replicate of EBs from three additional human lines of European ancestry to confirm that inter-species differences are not population-specific.

**Figure 1.**
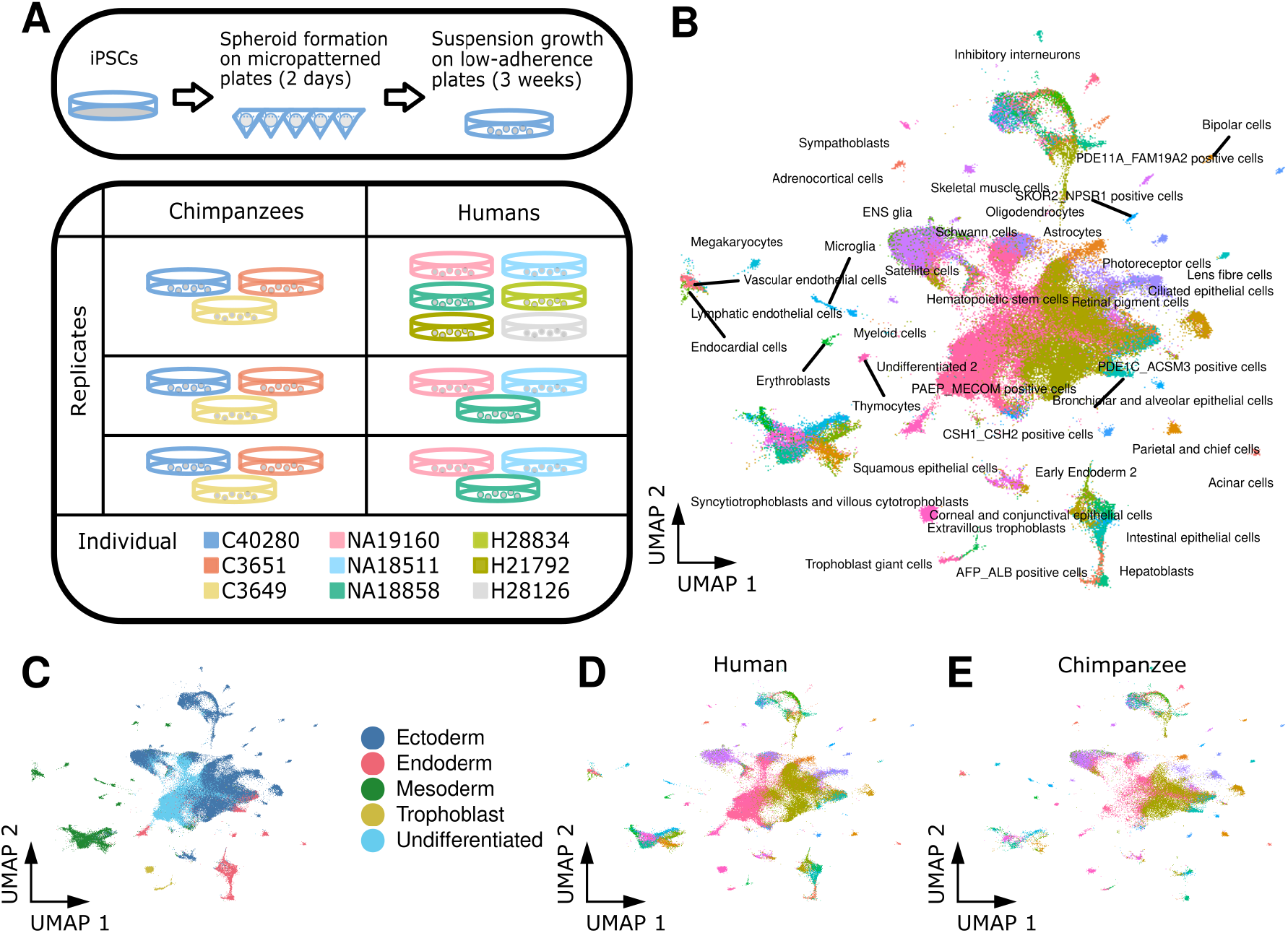
Human and chimpanzee embryoid bodies are composed of 86 cell types from all three germ layers. (**A**) Overview of experimental design. EBs were generated in three replicates of 6 to 9 human and chimpanzee iPSC lines. After two days in micropatterned plates, EBs were grown in low-adherence plates for 3 weeks before dissociating to single cells and sequencing with 10x scRNA-seq. (**B**) UMAP projection of all human and chimpanzee single cells after reference alignment. Cells are colored and labeled with the reference-assigned cell type. (**C**) The UMAP in (B) colored by germ layer of each cell type. The UMAP in (B) with only (**D**) human or (**E**) chimpanzee cells.

We formed EBs in micropatterned plates and cultured them for two days before transferring them to suspension culture. On day 21, we dissociated EBs into single-cell suspensions and collected single-cell transcriptomes using 10x Genomics scRNA-seq. To prevent systematic biases in transcriptomes due to differences in genome annotation, we aligned reads to a curated set of orthologous exons in the human and chimpanzee genomes. After filtering (see Materials and methods), we retained data from 115,269 cells (a median of 5,385 cells from each replicate) in which we detected the expression of a median of 3,149 genes (**Table S1**).

We first verified the presence of cells from the three germ layers in all samples by considering the expression of *SOX17, HAND1*, and *PAX6* as markers for endoderm, mesoderm, and ectoderm, respectively (**Figure S2**). In all samples, we also identified cells expressing *POU5F1* (Oct4), indicating the presence of undifferentiated cells (iPSCs). At low resolution, we identified 11 clusters of cells using Louvain clustering in Seurat (**Figure S2A**). In all clusters, we found cells from both species, with most clusters showing a nearly even proportion of human and chimpanzee cells (**Figure S2B and Table S2**).

To identify cell types within EBs, we considered human and chimpanzee gene expression data together with previously generated reference data containing cells from EBs, human embryonic stem cells (**hESCs**), and primary fetal tissues (Cao et al., 2020; Han et al., 2020; Rhodes et al., 2022). We labeled each of our cells based on its relative position within this reference (**Figure 1B**), which allowed us to partition cells into meaningful groups for subsequent analyses. In total, we identified 86 cell types present in both human and chimpanzee EBs (77 labels transferred from the reference and nine cell types identified *de novo*; see **Table S3**). These represent undifferentiated cells, cells from each of the three germ layers, and cells that closely resemble extra-embryonic cell types (**Figure 1C**). Cells from both species are well-distributed across all cell types (**Figure 1D-E**).

### Differential expression between humans and chimpanzees in 72 cell types

Next, we characterized differential expression (**DE**) between humans and chimpanzees in individual cell types. We excluded cell types that did not contain at least 5 cells in at least two replicates. We generated pseudobulk expression data for each of the remaining 72 cell types and performed DE analysis using a linear mixed model implemented in DREAM (Hoffman & Roussos, 2021). In addition to modeling the effect of species, we included random effects for replicate and individual. At a 5% false discovery rate (**FDR**), we identified DE genes in 70 cell types. Across all cell types, we identified 59,940 instances of DE genes between humans and chimpanzees; 10,457 (72% of all tested) genes are DE between the species in at least one cell type, with a range of two (chromaffin cells) to 3,712 (early ectoderm) DE genes in individual cell types (**Table S4 and Data S1**).

We explored the extent to which genes show inter-species DE in more than one cell type. To account for incomplete power to detect DE (given that we have only three individuals per species), we used Cormotif (Wei et al., 2015) to jointly analyze data from all 72 cell types simultaneously. This allowed us to share information across cell types, improving the power and accuracy of DE analysis. We identified 13 motifs of DE across cell types. At a posterior probability of 95%, we identified between 27 and 7,095 DE genes in each cell type, with a median of 2,718. 10,783 genes (74%) were DE in at least one cell type (**Table S4**). Compared to the underpowered single-tissue analysis, the mean number of cell types in which each gene was classified as DE increased from 4.1 to 14.7.

Examining the Cormotif results (**Figure 2A**), we found that the two largest classes of genes are those that are DE between species in all cell types (motif 11) and those that are not DE in any cell type (motif 1). These patterns of DE could indicate different degrees of selective constraint. Indeed, we found that the number of cell types in which a gene is classified as DE between species is significantly associated with coding sequence similarity between humans and chimpanzees (**Figure 2B**; *p* = 2.7 × 10^−6^). The ratio of non-synonymous to synonymous coding mutations across mammals (dN/dS) and the loss-of-function observed/expected upper bound fraction (**LOEUF**) (Karczewski et al., 2020) are also positively associated with the number of cell types in which a gene is classified as DE (**Figure 2C-D**; dN/dS *p* = 10^−13^; LOEUF *p* < 10^−15^). These observations are consistent with the notion that broad gene expression differences between humans and chimpanzees are generally associated with relaxation of evolutionary constraint. For a subset of broadly DE genes, it may be the case that directional selection has favored a new regulatory pattern. In support of this, we found that genes targeted by human accelerated regions (**HARs**) are also DE in a greater number of cell types (**Figure 2E**; *p* = 4.6 × 10^−5^).

**Figure 2.**
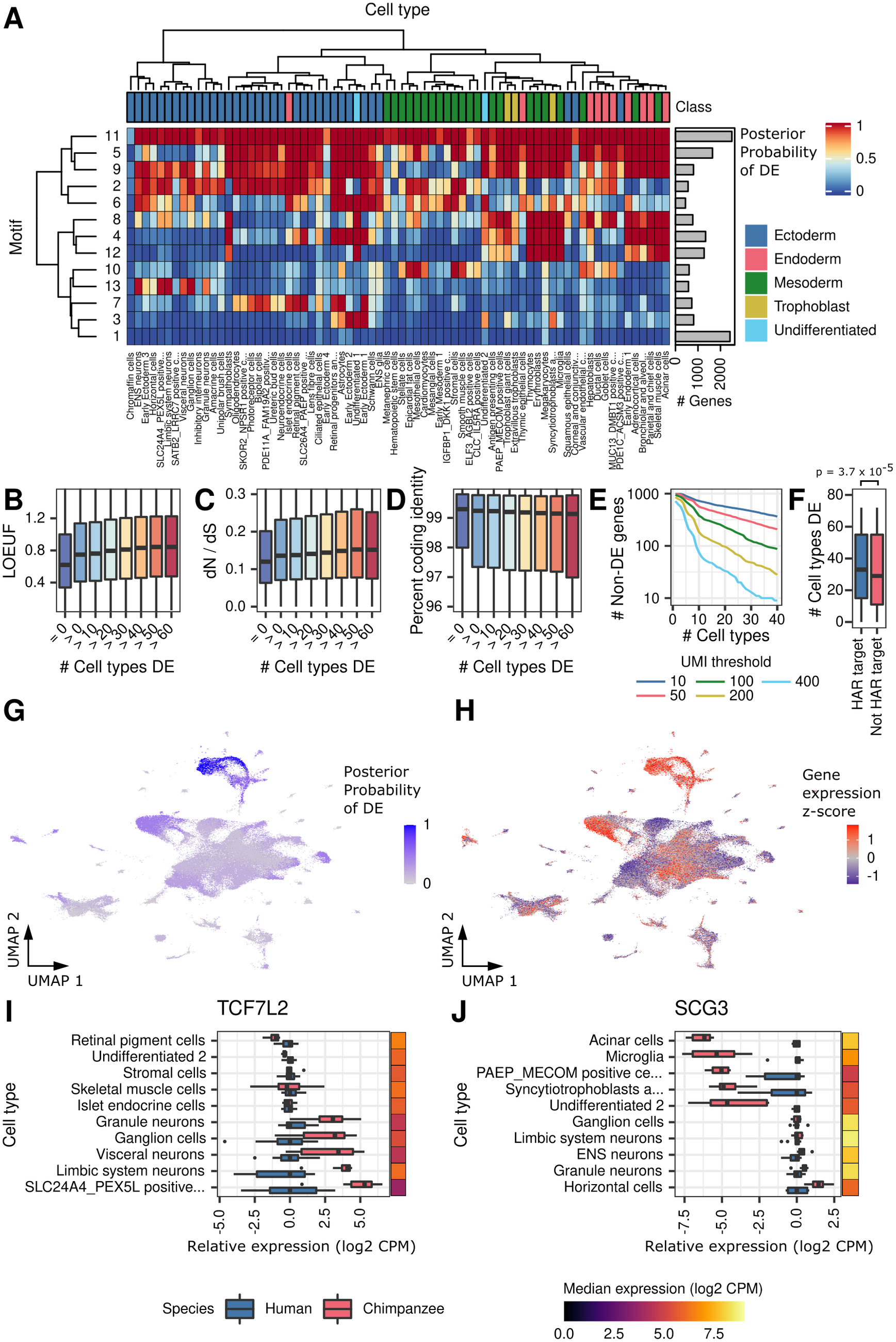
Sharing of differential expression across cell types. (**A**) Heatmap of correlation motifs across cell types. Cells are shaded according to the posterior probability of differential expression for each motif and cell type. Columns are clustered by mean gene expression. Inferred germ layers are given in **Table S3**. Boxplots of (**B**) Percent coding identity, (**C**) dN/dS, or (**D**) LOEUF in genes that are DE in zero or increasingly large numbers of cell types at posterior probability 0.5. (**E**) Boxplot showing the distribution of the number of cell types in which genes are DE for HAR targets and non-HAR target genes. (**F**) The number of non-DE genes remaining, requiring the specified absolute expression cutoffs in at least the given number of cell types. UMAP from **Figure 1B** is colored to show the (**G**) posterior probability of DE for genes in motif 13 or (**H**) the expression z-score of genes in motif 13. Boxplots show expression relative to humans in select cell types for (**I**) *TCF7L2* or (**J**) *SCG3*.

Genes with conserved expression levels across all cell types are of particular interest, as they may shed insight on core functions in both species. Yet, the identification of these genes is confounded by statistical power. To confidently classify these genes, we filtered non-DE genes by their absolute expression levels (**Figure 2F**). We classified hundreds of non-DE genes across all cell types, even at relatively stringent filters. For example, non-DE genes with at least 100 UMIs in at least 10 cell types in both species are enriched for core cellular processes, including protein transport and mRNA processing, splicing, and transport (**Table S5**).

Beyond genes that are DE in no cell types or all cell types, many of the remaining DE motifs have tissue-specific patterns. Inter-species DE in specific tissues could be driven by tissue-specific expression patterns, where genes may be DE everywhere they are expressed. In other cases, a gene may have acquired DE in a restricted set of cell types, despite being expressed in additional cell types. To examine these possibilities, we considered the expression levels of the genes in each motif across all cell types and compared them to patterns of DE (**Figure 2G-H and Figure S3**). We focused on motifs exhibiting tissue-specific DE while the genes are highly expressed, but are conserved in other cell types. We then identified genes with cell type-restricted acquisition of DE (see Materials and methods). We identified 3,906 genes (27%) that are broadly expressed but show cell-type-specific DE between humans and chimpanzees (**Data S2**).

Tissue-restricted DE could underlie important functional differences between humans and chimpanzees. For instance, the transcription factor TCF7L2 is a WNT modulator associated with human psychiatric and metabolic disorders (del Bosque-Plata et al., 2022, p. 7). *TCF7L2* is broadly expressed in EBs, but inter-species DE is restricted to neuronal subtypes, where it may underlie inter-species differences in neurodevelopment (**Figure 2I and Figure S4A-B**). Another example is *SCG3*, an obesity-associated gene (Tanabe et al., 2007) for which inter-species DE is restricted to a few cell types, including acinar (pancreatic) cells (**Figure 2J and Figure S4C-D**). This cell-type-specific pattern could underlie inter-species differences in digestive activity.

### Generation of EBs from tetraploid human chimpanzee hybrids

Next, we sought to further characterize mechanisms of inter-species DE. To quantify the contribution of *cis* versus *trans* regulatory divergence to inter-species differences in gene expression, we sequenced single EB cells from a human-chimpanzee fused stem cell line (Agoglia et al., 2021; Gokhman et al., 2021). We generated 65,247 fused single-cell transcriptomes from three replicates. Upon mapping these cells to the reference data, we observed cells from all 86 previously identified cell types (**Figure 3A**). We identified DE genes in each cell type using a paired Wilcoxon signed-rank test. At a 5% FDR, we observed DE in 67 cell types, with 13,842 genes classified as DE in at least one cell type (**Table S4**).

**Figure 3.**
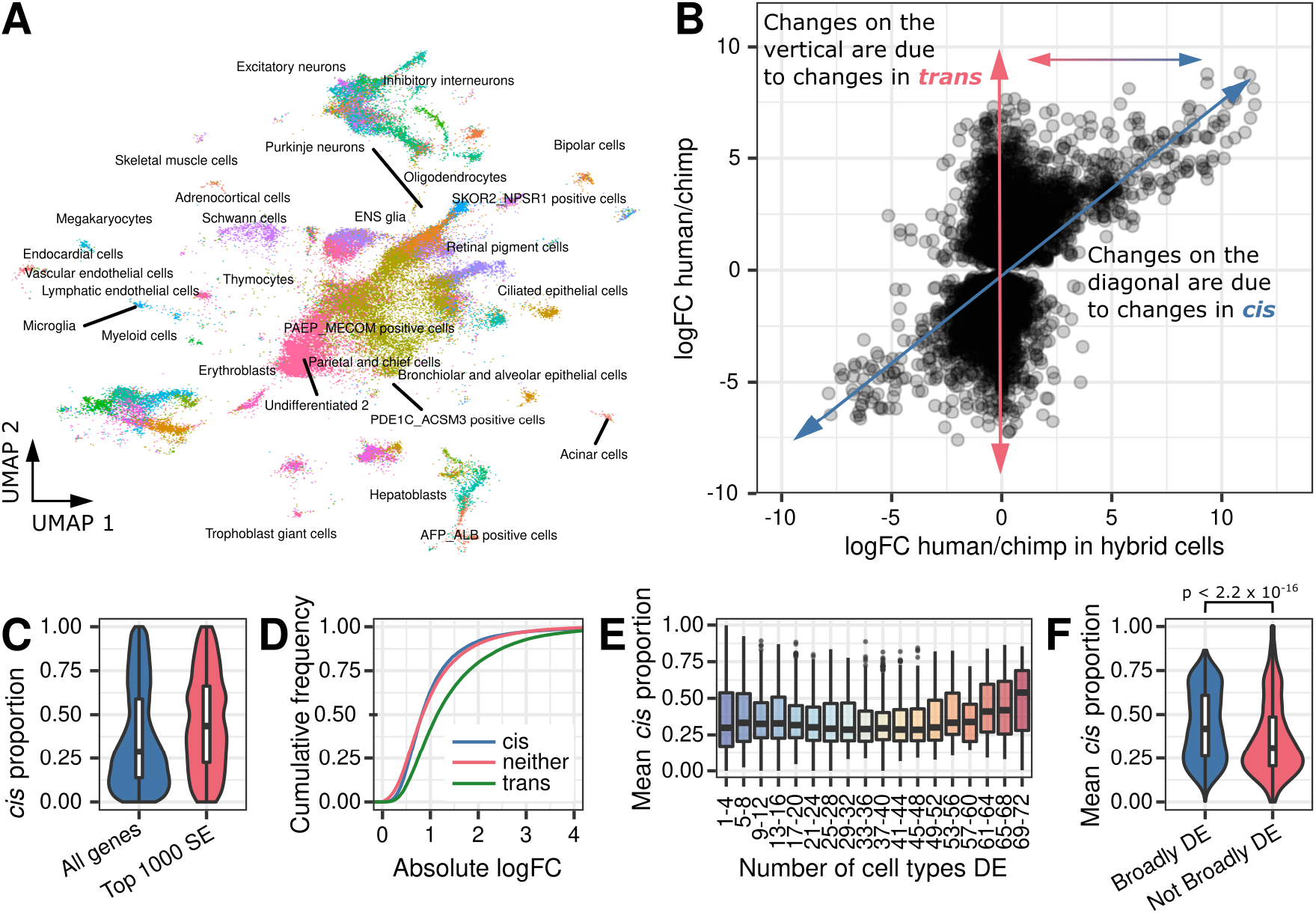
Allele-specific expression in tetraploid hybrid cells partitions differential expression into *cis* and *trans* components. (**A**) UMAP of hybrid cells after projection onto the space of **Figure 1B**. (**B**) The effect size in the hybrid vs. the effect size across humans and chimpanzees in DE genes (p < 0.001). (**C**) Boxplot of estimated *cis* proportion in the top 1,000 tests with the lowest estimate of standard error versus all other genes. (**D**) Empirical cumulative distributions of effect size (log2 fold-change from DREAM) for tests with high *cis* proportion (>0.75), low *cis* proportion (<0.25) or neither. (**E**) Boxplot of distribution of *cis* proportion for genes DE in increasingly large sets of cell types at posterior probability >95%. (**F**) Boxplot of *cis* proportion for genes DE in more than 80% of tested cell types (at posterior probability >95%) or all other genes.

Fused tetraploid cells contain the complete genome of both the human and chimpanzee parental lines; thus, genes from both species share the same *trans* environment. We estimated the proportion of gene expression divergence due to *cis* by comparing the change in gene expression observed between human and chimpanzee alleles in the tetraploid cells, to the change in gene expression observed between the original human and chimpanzee diploid cells (**Figure 3B**). Inter-species gene expression differences that cannot be explained by *cis* are assumed to be driven by changes in *trans* (though noise and measurement error will confound the *trans* estimate).

On average, we estimated that 70% of inter-species changes in gene expression are not due to changes in *trans* elements (**Figure 3C**), though as we pointed out, measurement error is expected to contribute to this estimate. Indeed, when we restrict this analysis to the 1,000 tests with the lowest error in the estimate of effect size, the proportion of *trans* regulation is decreased, as expected, to 62% (**Figure 3C**). We found that changes in *trans* elements are associated with larger effect sizes (**Figure 3D**; *p* < 10^−15^). We generally expect *trans* changes – particularly pleiotropic changes that affect multiple cell types – to be deleterious, and therefore, rare. However, a *trans* change that affects only one cell type is more likely to persist. Consistent with this expectation, we found that *trans* changes are depleted from genes that are broadly DE between the species – that is, the number of cell types in which a gene is classified as inter-species DE is negatively correlated with the proportion of DE that is explained by differences in *trans* (**Figure 3E-F**; *p* < 10^−15^).

We proceeded by arbitrarily classifying DE genes as *cis* if the mean estimated *cis* proportion across DE cell types was greater than 75%, or *trans* if the estimated *cis* proportion was less than 25%. We found that *cis* genes are associated with higher coding sequence divergence (**Figure 4A**; *p* = 2 × 10^−6^), higher dN/dS ratios (**Figure 4B**; *p* = 0.01), and higher LOEUF (**Figure 4C**; *p* =10^−15^) relative to *trans* genes. *Trans* genes are significantly enriched for functions related to development and morphogenesis (**Table S6**), and for many miRNA and transcription factor binding motifs (**Table S7**), the latter of which suggests a link between changes in expression and upstream regulators. For instance, *CLDN4* has a *trans* DE proportion of 93% and is a predicted target of the transcription factor *CEBPA*. Consistently, the pattern of DE across cell types for these two genes is highly correlated (**Figure 4D-E and Figure S5**; *ρ* = 0.88, *p* = 0.007).

**Fig 4.**
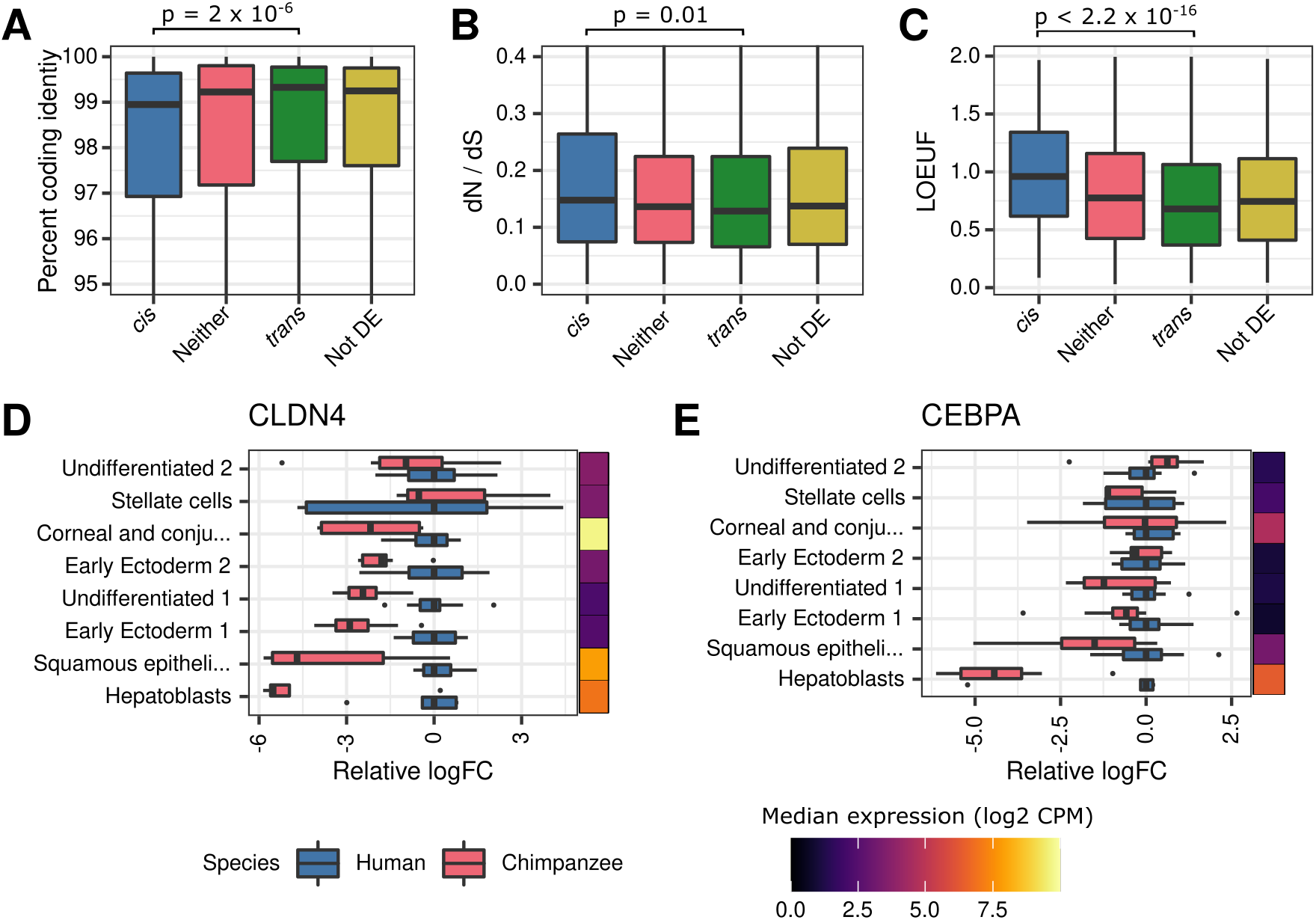
Genes with high *cis* proportion show evidence of relaxation of constraint. Boxplot showing the distribution of (**A**) percent coding identity (**B**) dN/dS or (**C**) LOEUF for genes with a high, moderate, or low *cis* proportion. Boxplots of showing expression relative to mean human expression in select cell types for (**D**) *CLDN4* or (**E**) *CEPBA*.

## Discussion

We used EBs to create the most comprehensive comparative catalog of gene expression levels in humans and chimpanzees to date, including single-cell data from 86 developing and mature cell types across multiple germ layers and tissues. Importantly, the EB model system allowed us to generate each of these cell types from every human and chimpanzee individual using uniform processing, such that we did not confound cell type with individual, species, or processing method in our analysis. Our balanced study design allowed us to explore gene regulatory divergence at an unprecedentedly high cellular and temporal resolution.

We confirmed that gene regulatory divergence is ubiquitous across cell types, even in closely related species. More than two thirds of tested genes are DE in at least one cell type, and in the majority of cell types there are hundreds to thousands of DE genes between the species. We identified thousands of genes that are DE in a subset of the cell types in which these are expressed, which suggests that there are quite a few functional changes in tissue-specific gene regulation between the two species.

We were somewhat surprised to find a large subset of genes whose expression is conserved in all of the cell types we were able to study. We did not expect such a large fraction of genes to be regulated in a similar way in humans and chimpanzees in over 70 different cell types, as this requires that the regulation of these genes is of sufficient functional importance in all of these cell types to evolve under relatively strong constraint. It is of course possible that with data from more individuals, we would find that a subset of the ‘generally conserved’ genes are in fact DE between the species in some of the cell types; nevertheless, this observation is intriguing and suggests that we should reconsider our assumptions regarding the degree of context-specific gene regulation.

Turning our attention to regulatory mechanisms, the use of fused cell lines allowed us to assess the relative contribution of *cis* and *trans* changes to regulatory evolution – an approach that has not been feasible in primates until recently (Agoglia et al., 2021; Gokhman et al., 2021; Starr et al., 2022). We found that *cis* changes are common and can impact any number of cell types, as expected given the potential for spatial and temporal specificity of *cis* elements. Our observation that *trans* changes are associated with inter-species DE in fewer cell types than *cis* changes may seem counterintuitive, because one expects *cis* changes to be more specific than changes in *trans*. However, *trans* changes are more likely to be associated with pleiotropic deleterious outcomes than changes in *cis*. Thus, while *trans* changes are likely more exposed to negative selection, they may persist if their impact is restricted to a small number of cell types and thus drive the observed correlation between breadth of DE and proportion of *trans*.

We also found that fused lines help to identify putative regulatory relationships between genes. By combining correlated patterns of DE across cell types with binding motif enrichment in the promoters of *trans*-regulated DE genes, we demonstrated that *CEBPA* may drive tissue-specific differential expression of *CLDN4*. In principle, this approach could be used to identify key sets of *cis*-regulated genes that drive divergence in expression between humans and chimpanzees. Future studies of human-chimpanzee hybrid EBs could also explore how *cis* and *trans* differences are distributed through gene regulatory networks.

Collectively, our work validates EBs as a powerful model system for comparative studies of gene expression in humans and chimpanzees. The EB system confers several advantages over existing models. First, the use of EBs sidesteps the many pitfalls of working with frozen tissues, which have severely limited the scope of comparative studies in primates. Frozen tissues from non-human apes must be collected opportunistically, making it difficult to sample broadly across tissues, obtain samples from more than a handful of individuals per species, and design unbiased studies that are balanced with respect to species, tissue, and individual (Blake et al., 2020). Moreover, frozen tissues cannot be used to study how gene regulation changes over time or in response to external perturbations. While iPSC-derived cell types and organoids provide a way to explore gene regulatory dynamics *in vitro* (Agoglia et al., 2021; Elorbany et al., 2022; Pollen et al., 2019; Shibata et al., 2012; Ward & Gilad, 2019), EBs can be used to efficiently generate multiple cell types with a fraction of the effort it takes to differentiate a single iPSC line. Indeed, EBs permit the study of gene regulation across dozens of cell types, individuals, developmental time points, and environmental exposures, which will allow future studies to explore gene regulatory divergence in apes in greater detail than has been possible thus far.

In particular, gene expression response phenotypes have not been widely explored in apes, even using *in vitro* models. Understanding how human and chimpanzee cells respond to external perturbations can reveal how gene regulatory divergence affects transcriptional responses that are relevant to human-specific traits and diseases, and may reveal functionally important genes and pathways that affect fitness. In a previous study, we examined the immune response to infection in primary monocytes from humans, chimpanzees, and rhesus macaques (Barreiro et al., 2010). We observed that while the regulatory response to bacterial infection is largely conserved between species, the regulatory response to viral infection is often lineage-specific. For example, we found chimpanzee-specific immune signaling pathways to be enriched for HIV-interacting genes, which could explain why HIV-infected chimpanzees exhibit relatively strong resistance to AIDS progression (Barreiro et al., 2010).

More recently, we used iPSC-derived cardiomyocytes from humans and chimpanzees to examine inter-species differences in response to hypoxia, a symptom of myocardial ischemia (Ward & Gilad, 2019). Despite differences in myocardial ischemia risk, heart development, and other cardiovascular traits between the two species, we observed that the response of human and chimpanzee cardiomyocytes to hypoxia was largely conserved. We also found that hypoxia-induced transcription factors bind more frequently to conserved hypoxia response genes than to chimpanzee-specific hypoxia response genes, suggesting that changes to transcription factors or their binding sites might underlie inter-species differences in hypoxia (and possibly, differences in myocardial infarction risk). In each of these studies, we identified functionally important genes and mechanisms by comparing the transcriptional responses of a single cell type to a disease-relevant exposure. The EB model system makes it possible to perform similar studies in dozens of cell types at once, including transient, developmental cell types that are not accessible from adult tissues.

In summary, EBs are a powerful *in vitro* model system that can be used to address a number of outstanding questions in primate comparative genomics. Our initial exploration of human and chimpanzee EBs resulted in the first comparative database of *cis* and *trans* regulatory divergence between the species, which includes gene expression data from more than 70 human and chimpanzee cell types. Future studies will take full advantage of this model by including additional human and chimpanzee cell lines, by studying how *cis* and *trans* changes interact to drive adaptation and divergence, and by exploring divergent transcriptional responses to biomedically relevant perturbations between the two species.

## Materials and methods

### Samples

In this study we included six human and three chimpanzee iPSC lines. The three human lines were derived from unrelated individuals from the Yoruba population in Ibadan, Nigeria (YRI) as part of the International HapMap Collection. These lines were reprogrammed from lymphoblastoid cell lines and characterized in (Banovich et al., 2018). Two of these lines are female (18511, 18858) and one is male (19160). We included three additional fibroblast-derived human iPSC cell lines characterized in (Gallego Romero et al., 2015). Two of these lines are female (H28834, H21792) and one is male (H28126). We included three fibroblast-derived chimpanzee iPSC lines, also characterized in (Gallego Romero et al., 2015). These also included two females (C40280, C3651) and one male (C3649). We obtained the tetraploid human/chimpanzee hybrid iPSC line (HLI-25) from Dr. Hunter Fraser.

### iPSC maintenance

We maintained feeder-free iPSC cultures on Matrigel Growth Factor Reduced Matrix (CB-40230, Thermo Fisher Scientific) with StemFlex Medium (A3349401, Thermo Fisher Scientific) and Penicillin/Streptomycin (30002Cl, Corning). We grew cells in an incubator at 37°C, 5% CO2, and atmospheric O2. Every 3-5 days thereafter, we passaged cells to a new dish using a dissociation reagent (0.5 mM EDTA, 300 mM NaCl in PBS) and seeded cells with ROCK inhibitor Y-27632 (ab120129, Abcam).

### Embryoid body formation and maintenance

We formed EBs using the STEMCELL AggreWell™400 protocol according to the manufacturer’s directions. Briefly, we generated a single-cell suspension of iPSCs by incubating them for 5 to 13 minutes with 300 mM NaCl in PBS, followed by gentle pipetting. We seeded iPSCs into an AggreWell™400 24-well plate at a density of 1,000 cells per microwell. We cultured the aggregates in AggreWell™ EB Formation Medium (05893, STEMCELL) with ROCK inhibitor Y-27632 for 24 hours. After 24 hours, we replaced 1mL of the media with fresh EB Formation Medium. After another 24 hours, we harvested the EBs and placed them on an ultra-low attachment 6-well plate (CLS3471-24EA, Sigma) in E6 media (A1516401, ThermoFisher Scientific). We replaced the media every other day for the next 19 days, for a total of 21 days of EB culture.

We formed EBs on three separate days. The first day we included all six human lines and all three chimpanzee lines. For the next two experimental replicates, we included all three chimpanzee lines, but only the three YRI human lines.

### Embryoid body dissociation

On day 21 post-formation, we dissociated EBs into a single-cell suspension. We washed the cells in PBS (Corning 21-040-CV) and then incubated them in 1mL of AccuMax dissociation reagent (STEMCELL 7921) for 10 minutes at 37°C. After 10 minutes, we gently pipetted the EBs for 30 seconds using a clipped p1000 tip. We repeated this every five minutes until we achieved a single-cell suspension, up to a maximum of 35 minutes. Next, we stopped dissociation by adding 5mL of ice-cold E6 media. We then filtered cells using a 40um strainer (Fisherbrand 22-363-547). We washed the cells three times using ice-cold PBS + 0.04% BSA. Prior to running the cells on the 10X Chromium controller, we mixed the cells together, including an equal number of cells from each individual.

### Library preparation and sequencing

We generated scRNA-seq libraries using the 10X Genomics 3’ scRNA-seq v3.0 kit. We targeted 10,000 cells per lane of the 10X chip. In the first replicate, with nine individuals, we initially collected nine lanes (∼90,000 cells). For the remaining two replicates we collected four lanes (∼40,000 cells). We performed all sequencing using 100bp paired-end sequencing on a HiSeq 4000 at the University of Chicago Functional Genomics Core Facility. We initially pooled and replicated libraries 1-2 and 6-9 and sequenced on one lane, and pooled libraries 1 and 3-5 on another lane. Our preliminary analysis indicated that library 2 was poor quality and we excluded it from further analyses. Libraries 3, 4, and 5 contained fewer cells than expected, but otherwise appeared to be good quality. We re-pooled the eight remaining libraries using half the amount of libraries 3-5 to account for lower library complexity. We sequenced this pool on eight lanes of the HiSeq 4000. We also pooled and sequenced libraries from replicates 2 and 3 on one lane each of the HiSeq 4000. After preliminary analysis, we performed an additional six and four lanes of sequencing on the HiSeq400 for replicate 2 and replicate 3 libraries, respectively. For each replicate, the number of sequencing lanes was chosen such that we obtained 50% saturation as assessed by 10X cellranger software.

### Orthologous exon reference generation

In order to make quantitative comparisons across species that are not biased by differences in mapping annotation quality across species, we generated a database of orthologous exons. We followed the general approach of (Blekhman et al., 2010; Pavlovic et al., 2018) but updated this database to use the most recent version of the chimpanzee genome assembly (Clint_PTRv2/panTro6). Briefly, we mapped the human genome annotation (hg38/Ensembl98) onto the chimpanzee genome using three separate BLAT steps. First, we used BLAT on every protein-coding human exon to identify the ortholog in chimpanzee. We kept exons that had at least 80% sequence identity. For exons with multiple locations at greater than 80% sequence identity, we retained the exon that shared the most neighboring exons (within 100 kb) with the original reference. We also removed exons with indels greater than 20% of the exon length. Next, we used BLAT to map these chimpanzee exons back to the human genome. We retained exons mapped back to their original location in human. Finally, we used BLAT to the map the chimpanzee exons to the chimpanzee genome. We retained exons that map back to the original location in chimpanzee. The resulting reference contains 1,077,831 orthologous exons from 19,787 genes.

### Alignment, species assignment, demultiplexing, filters and normalization of human and chimpanzee libraries

We compiled three separate references using the cellranger *mkref* command from 10x Genomics: one human, one chimpanzee, and one combined human and chimpanzee reference. Each reference was generated using the appropriate species annotation from the orthologous exon database described above. We used cellranger count to map reads to each of these three references. We used the combined genome reference to assign species to each barcode. We assigned a cell barcode to a species if greater than 90% of uniquely mapping reads mapped to that species’ genome.

We used demuxlet to assign each droplet to an individual (Kang et al., 2018). We used genotypes for NA19160, NA18858 and NA18511 from the 1000 genomes project (Consortium, 2012). Genotypes for human and chimpanzee individuals H21792, H28126, C3649, C3651, and C40280 were obtained from Dr. Gregory Wray. We generated genotype calls for the remaining individual, H28834, using RNA-sequencing data from (Blake et al., 2018) and the GATK best-practices pipeline for calling genotypes from sequencing data.

We filtered the data to remove droplets that had fewer than 1,000 genes detected. We also removed droplets where mitochondrial reads represented more than 20% of total reads or fewer than 0.1% of total reads. We processed the data using the SCTransform pipeline from Seurat (Hafemeister & Satija, 2019) using 5,000 variable genes and obtained Pearson residuals for all genes.

### Reference integration and cell type assignment of human and chimpanzee cells

We have previously reported a joint reference containing cells from human EBs (Rhodes et al., 2021) as well as cells from a fetal cell atlas (Cao et al., 2020) and human embryonic stem cells (**hESCs**) (Han et al., 2020). We mapped the human and chimpanzee cells onto this reference. Briefly, we embedded the human and chimpanzee cells in the PCA space of the joint reference. Next, we used Harmony (Korsunsky et al., 2019) to correct the PCs of the human and chimpanzee datasets while holding the reference PCs fixed.

In order to assign cell type identities from the fetal cell reference, we first assigned cluster midpoints (by taking the mean of each harmony-corrected PC) for each cell type in the fetal cell atlas as well as hESCs. We then assigned a cell identity to each cell in the human and chimpanzee EB dataset by finding the closest reference cluster midpoint by Euclidean distance. This resulted in a substantial number of cells that were assigned to hESCs, meaning they more closely resembled undifferentiated cells than anything within the fetal reference. These cells represent both undifferentiated cells and immature or transient developing cell types not present in either reference. To assign identities to these cells, we performed unsupervised clustering using the Louvain algorithm (in Seurat, using resolution 0.1). This resulted in 9 new cell types. We then reassigned every human and chimpanzee EB cell using the fetal cell reference midpoints, as well as midpoints for each of these 9 *de novo* cell types.

### Alignment, filters, normalization, and cell type assignment of hybrid libraries

We performed two separate alignments of the tetraploid human-chimpanzee hybrid cell lines using cellranger count. We aligned all of the data to the standard human reference (2020-A, obtained from 10x Genomics) as well as the combined orthologous exon reference described above. We used the human-aligned data to filter, normalize, integrate with the reference, and assign cell identities. We removed droplets that had fewer than 2,000 genes detected, as well as droplets with fewer than 0.1% or more than 20% of reads aligning to mitochondrial genes. We processed the data using SCTransform with 5,000 variable genes, returning residuals for all genes. Using the procedure described above, we then aligned our data with the joint reference, including all human and chimpanzee data generated in this study. We assigned each hybrid cell the identity of the nearest reference cell by Euclidean distance in Harmony-corrected PC space.

We used data aligned to the combined human and chimpanzee reference for all quantitative comparisons. By default, cellranger discards sequencing reads that do not uniquely map to the genome. Therefore, reads that remain after alignment represent fragments that can be uniquely assigned to the allele of one species. We retained droplets that passed filters in the human-aligned data. We also transferred relevant metadata, including cell type identities, from the human-aligned data.

### *Differential expression between humans and chimpanzees with* DREAM

In order to perform differential expression (**DE**) analysis, we generated pseudo-bulk expression by taking the sum of counts for all cells that came from the same individual, replicate, and cell type. We than removed any samples with 5 or fewer cells. For each cell type, we filtered genes using the *filterByExpr* function from the R package edgeR(Robinson et al., 2010), setting the variable *min*.*count* to 5. We performed DE analysis using DREAM (Hoffman & Roussos, 2021) separately in each cell type, setting species as a fixed effect and individual and replicate as random effects. We estimated size factors using trimmed mean of M-values (**TMM**) with singleton pairing and we performed cyclic loess normalization.

### *Joint differential expression with* Cormotif

The package Cormotif (Wei et al., 2015) uses *t*-statistics from LIMMA (Smyth, 2004), performed in multiple conditions, to share information and improve power of DE analysis. The software takes normalized gene expression measurements as input. This may be inappropriate for single-cell RNA-sequencing projects, where the number of cells per sample may vary by more than an order of magnitude. We modified the Cormotif package to accept UMI counts as input. To generate *t*-statistics, our modified package runs limma-voom (Law et al., 2014) in order to apply precision weights to each sample. We used the same gene filters as we used for DREAM. We also used TMM and cyclic loess normalizations. The modified Cormotif package that implements this is available on github at https://github.com/kennethabarr/CormotifCounts.

### Identification of cell type restricted differential expression

To visualize the expression patterns of genes within correlation motifs, we generated a motif score for each motif in each cell. For each motif, we first took the product of the SCTtransform residuals for each gene and the probability that each gene is a member of that motif. We then summed these motif membership-weighted SCTransform residuals for all genes. We then computed z-scores of this metric to visualize distributions of motif expression.

To identify single genes with tissue-restricted patterns of DE, we first filtered for genes with greater than 95% posterior probability of DE in at least one cell type. Within this set, we identified genes that had posterior of probability of DE of less than 5% in at least one cell type. To ensure that differential power could not explain the lack of DE in the conserved cell type, we required that the conserved cell type have at least 100 total observed UMIs for that gene in both humans and chimpanzees.

### Differential expression in tetraploid hybrid cells

For every gene and cell type, we performed a paired Wilcoxon signed rank test to compare the within-cell difference between counts of reads assigned to human alleles vs chimpanzee alleles. We used the row_wilcoxon_paired function from the matrixTests package to implement this (Koncevičius, 2021). We then adjusted p-values within each cell type using the Benjamini-Hochberg procedure (Benjamini & Hochberg, 1995).

### *Estimates of* cis *proportions*

To estimate the relative contribution of allele-specific changes (*cis* changes) to human and chimpanzee gene expression divergence, we compared the effect size between humans and chimpanzees to the effect size within tetraploid hybrid cells where the *trans* environment is fixed. We used the procedure outlined in (Agoglia et al., 2021; Gokhman et al., 2021). Briefly, we compute the *cis* component as the absolute value of the log2 fold change between humans and chimpanzees measured in hybrid cells, which we denote abs(hybrid.logFC). The *trans* and non-genetic component is the absolute value log2 fold change between humans and chimpanzees measured in human and chimpanzee cells, minus the *cis* component. For this we used the effect size estimate computed by DREAM. We denote the *trans* and non-genetic component as abs(dream.logFC – hybrid.logFC). Thus, the *cis* proportion is given by abs(hybrid.logFC) / (abs(hybrid.logFC) + abs(dream.logFC – hybrid.logFC)).

### External data sources

We obtained percent coding identity between humans and chimpanzees from BioMart (Kinsella et al., 2011). Mammalian dN/dS was obtained from a study of 29 mammalian genomes (Lindblad-Toh et al., 2011). LOEUF was obtained from a study of 141,456 human genomes (Karczewski et al., 2020). Human accelerated regions and their targets were obtained from a recently compiled set of 3,171 regions (Girskis et al., 2021).

### Gene ontology and gene set enrichment

We performed gene ontology analysis using the R package topGO (Alexa & Rahnenfuhrer, 2022). We used the Fisher exact test as the statistic, and set the nodeSize parameter to 10. We performed gene set enrichments using the package *fgsea* (Korotkevich et al., 2021), with the eps parameter set to 0. We obtained gene sets from the Molecular Signature Database (MSigDB v7.5.1) (Liberzon et al., 2011). For gene set enrichment of genes generally regulated in *trans*, we ranked genes based on the mean *trans* proportion across all cell types in which each gene was DE. We provided this ranking as input to *fgsea* and used the positive score type.

### Statistics

For all correlations reported in the main text, we computed p-values using Spearman’s rank-order correlation test, implemented in R with the function corr.test with option method=‘s’. For all comparisons of the difference between distributions, we computed p-values using the two-sided Mann–Whitney *U* test, computed in R with the function wilcox.test.

## Acknowledgments

We thank all members of the Gilad lab for helpful discussions, the Functional Genomics Core Facility at the University of Chicago for sequencing the RNA-seq libraries, and Natalia Gonzales for editing the manuscript.

## Author contributions

Conceptualization: KAB, KLR, YG

Methodology: KAB, KLR

Software: KAB

Validation: KLR, KAB

Formal analysis: KAB

Investigation: KAB, KLR

Resources: YG

Data curation: KLB

Writing – original draft: KAB, YG

Writing – review & editing: KAB, YG

Visualization: KAB

Funding acquisition: YG

Project administration: YG

Supervision: YG

## Data and materials availability

All raw and processed sequencing data generated in this study have been submitted to NCBI’s Gene Expression Omnibus (GSE201516). Code generated for this project is available on GitHub (https://github.com/kennethabarr/CormotifCounts and https://github.com/kennethabarr/HumanChimp).

## Supplementary Information and Extended Data

### Figs. S1 to S5

**Figure S1.**
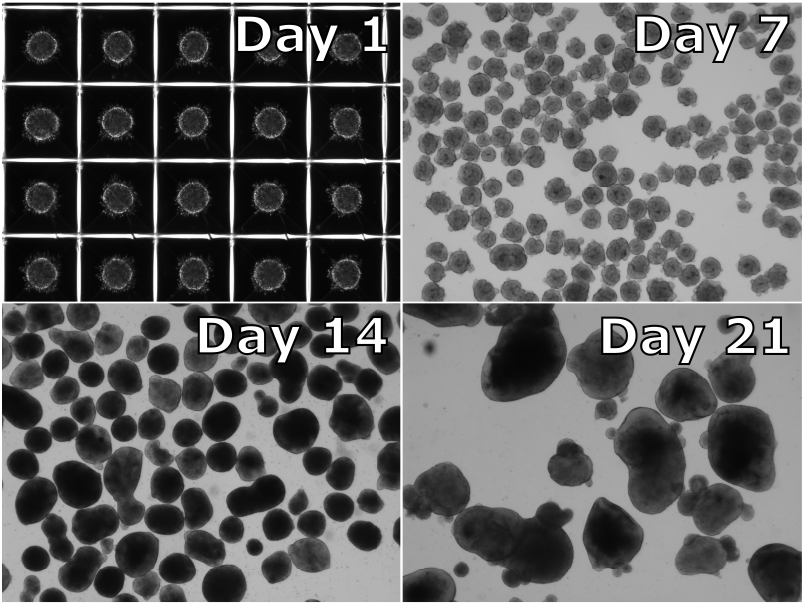
Representative images of embryoid bodies. Images of EBs at days 1, 7, 14, and 21 post-formation. Images are taken on an EVOS 2 FL at 4x resolution.

**Figure S2.**
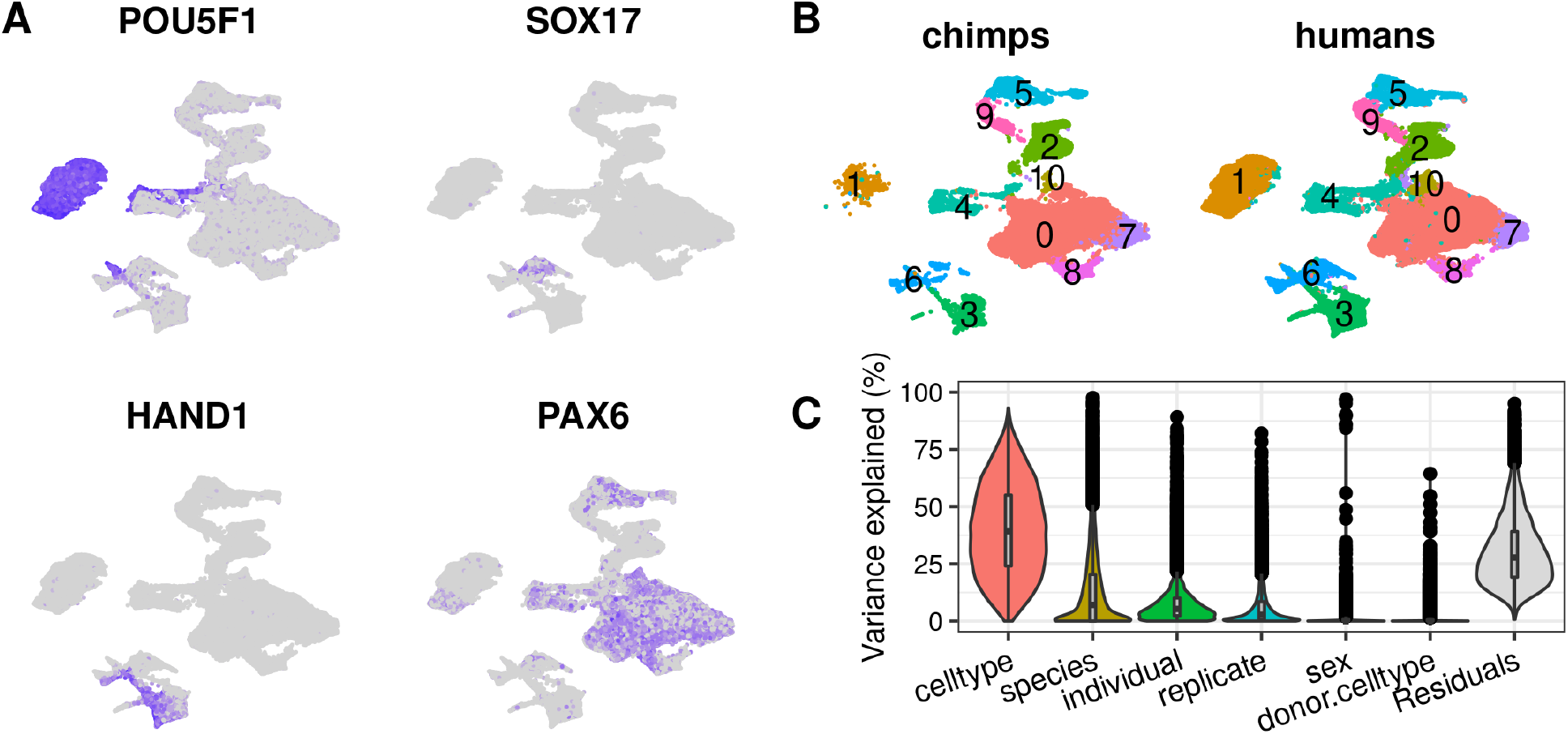
Cell type composition in embryoid bodies without external reference integration. (**A**) UMAP of EB cells highlighting expression of germ layer markers *POU5F1, SOX17, HAND1*, and *PAX6*. (**B**) UMAP split by species. Cells are colored by Seurat clusters at resolution 0.1. (**C**) The percent of variance explained by biological and technical factors using the VariancePartition R package.

**Figure S3.**
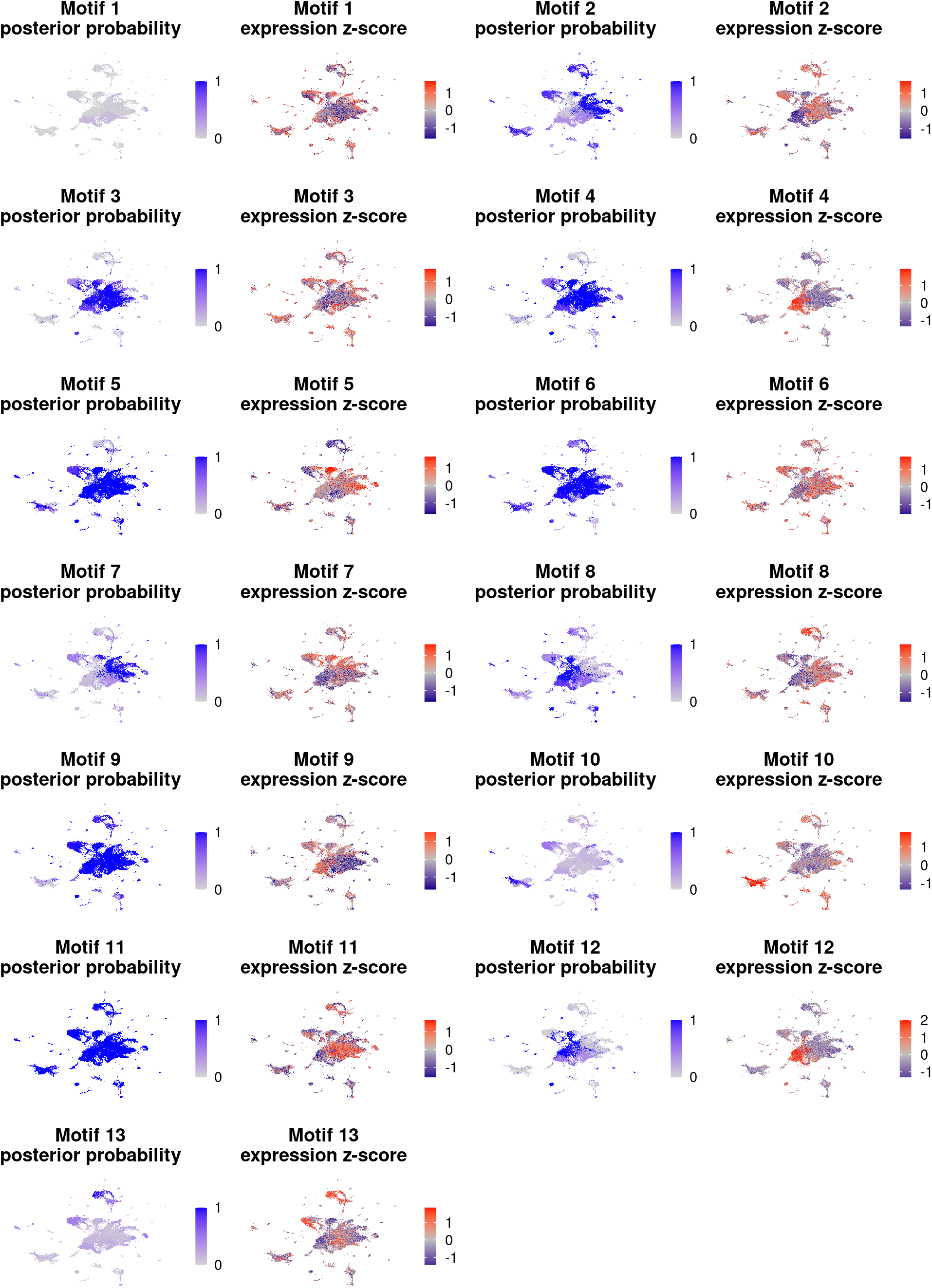
UMAPs of Cormotif posterior probability of differential expression and gene expression z-scores. UMAPs from **Figure 1B** with cells colored according to their posterior probability of DE or gene expression z-score (see Materials and methods).

**Figure S4.**
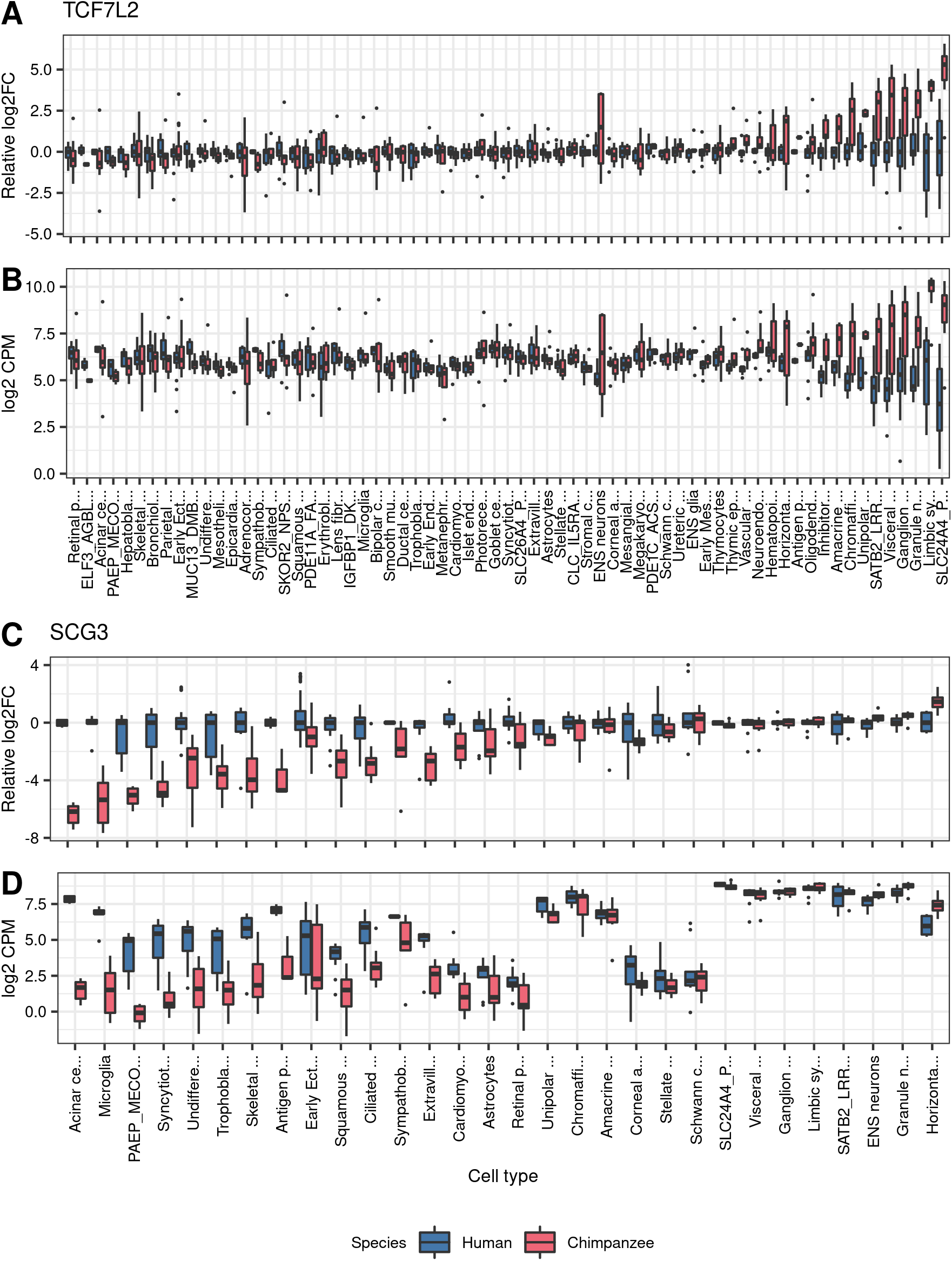
Boxplots of *TCF7L2* and *SCG3* expression. (**A**) Expression of *TCF7L2* in all tested cell types, relative to the mean expression in humans. (**B**) Expression of *TCF7L2* in all tested cell types. (**C**) Expression of *SCG3* in all tested cell types, relative to the mean expression in humans. (**D**) Expression of *SCG3* in all tested cell types.

**Figure S5.**
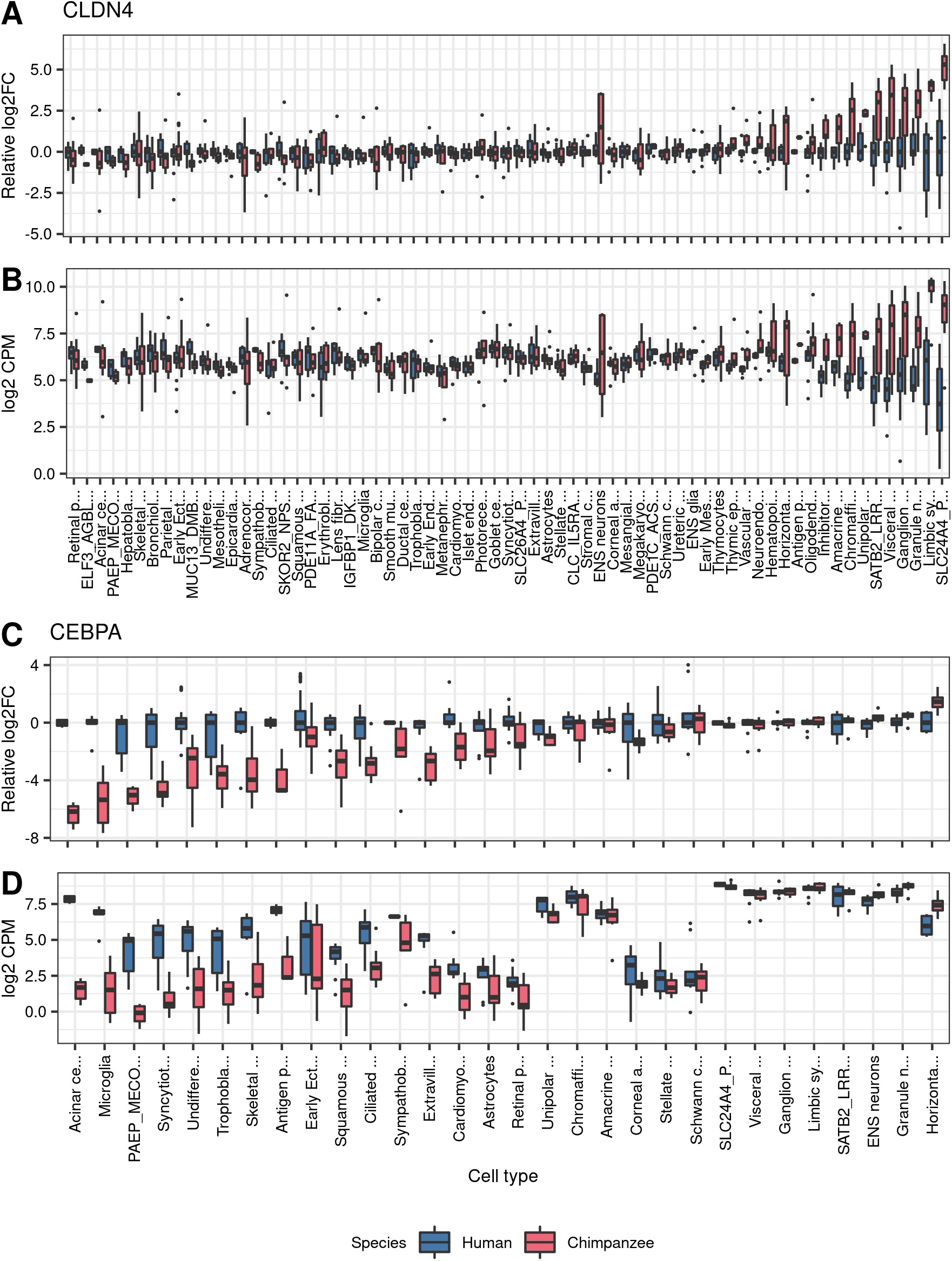
Boxplots of *CLDN4* and *CEBPA* expression. (**A**) Expression of *CLDN4* in all tested cell types, relative to the mean expression in humans. (**B**) Expression of *CLDN4* in all tested cell types. (**C**) Expression of *CEBPA* in all tested cell types, relative to the mean expression in humans. (**D**) Expression of *CEBPA* in all tested cell types.

### Tables S1 to S7

**Table S1.** Number of cells from each individual and replicate after quality control filters.

|  | Replicate1 | Replicate2 | Replicate3 |
| --- | --- | --- | --- |
| NA19160 | 7921 | 4937 | 7186 |
| NA18511 | 5383 | 5150 | 6068 |
| NA18858 | 6805 | 5099 | 5385 |
| H28834 | 5801 |  |  |
| H21792 | 5851 |  |  |
| H28126 | 5962 |  |  |
| C40280 | 3737 | 3569 | 6841 |
| C3651 | 5586 | 4615 | 4652 |
| C3649 | 5263 | 3560 | 5898 |

**Table S2.** Number of cells from each species in each cluster without reference integration. Clusters are those defined by Seurat at resolution 0.1, and are illustrated in **Figure S2B**.

| cluster | human | chimp |
| --- | --- | --- |
| 0 | 20921 | 24653 |
| 1 | 20501 | 398 |
| 2 | 4650 | 5931 |
| 3 | 7061 | 910 |
| 4 | 4106 | 2554 |
| 5 | 2119 | 3663 |
| 6 | 4702 | 379 |
| 7 | 3592 | 1196 |
| 8 | 653 | 2742 |
| 9 | 1704 | 1102 |
| 10 | 1539 | 193 |

**Table S3.**
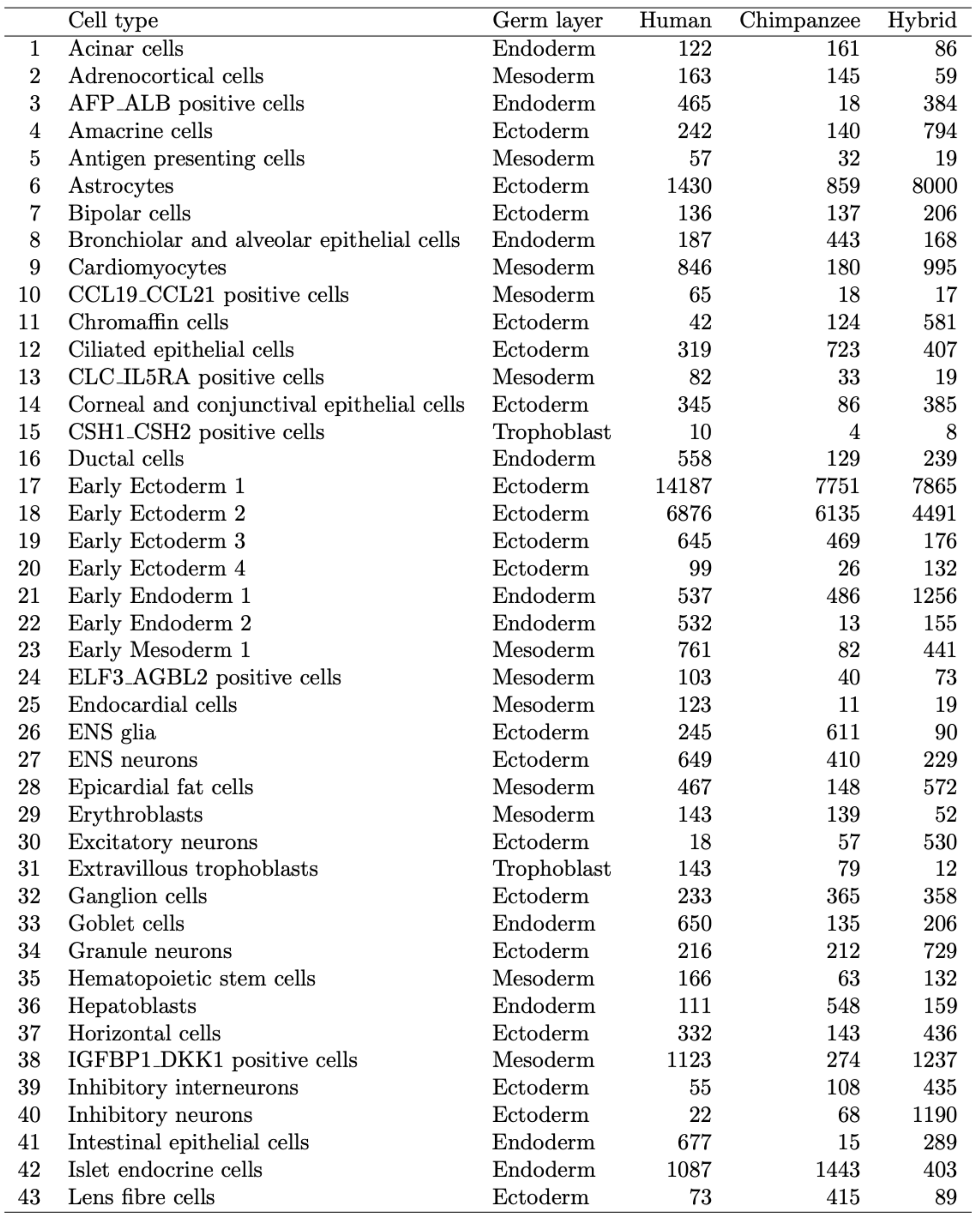
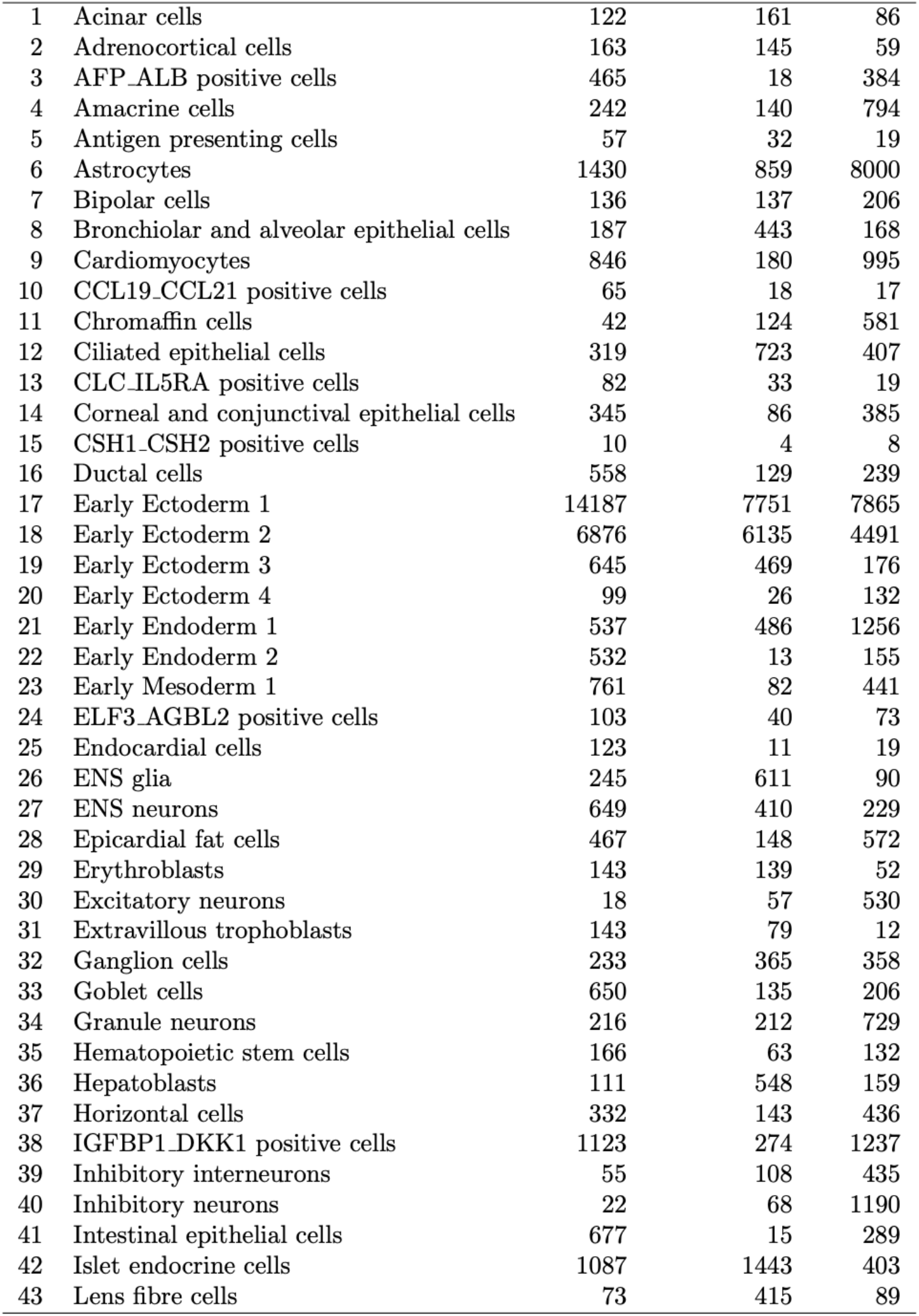
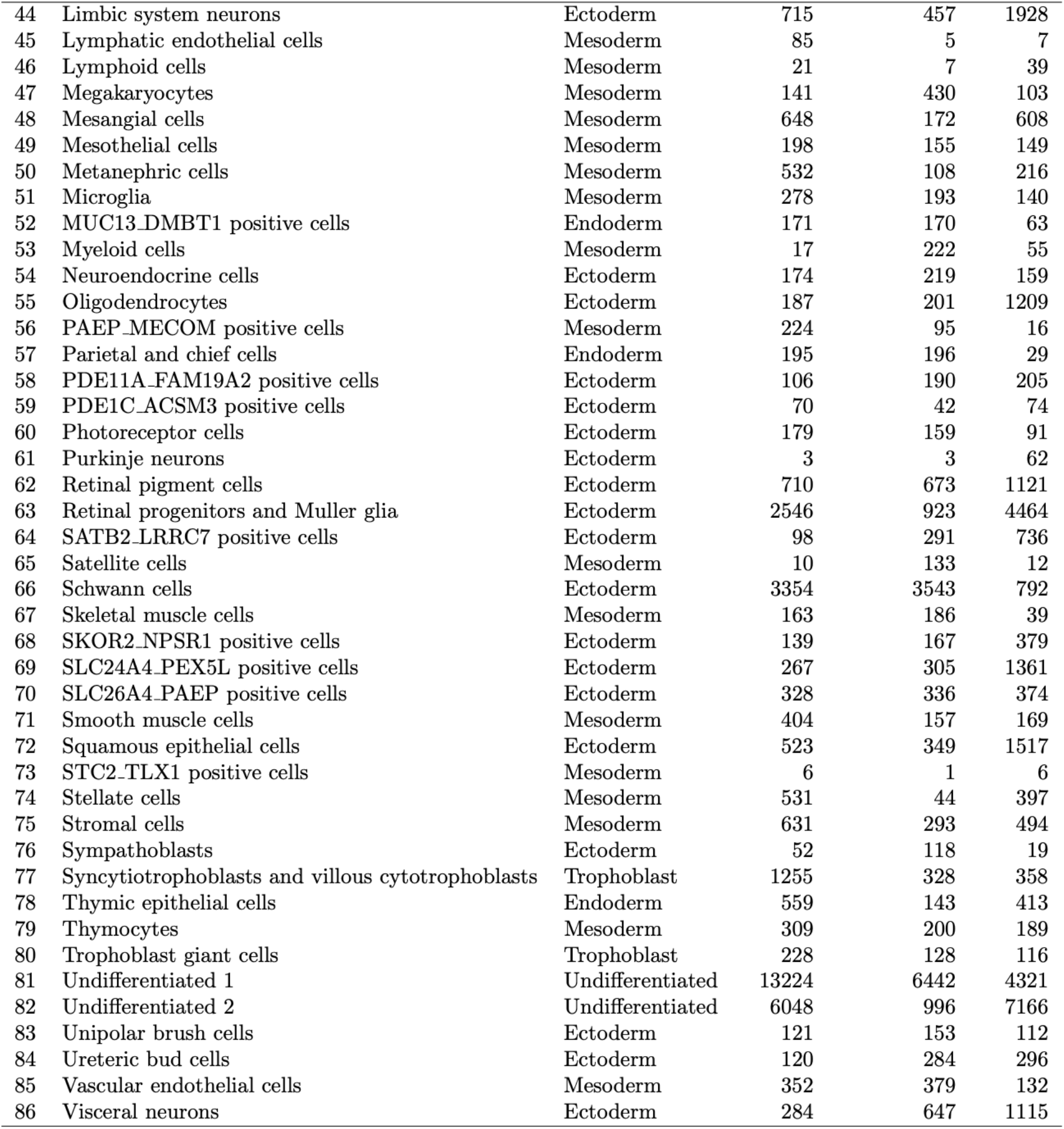
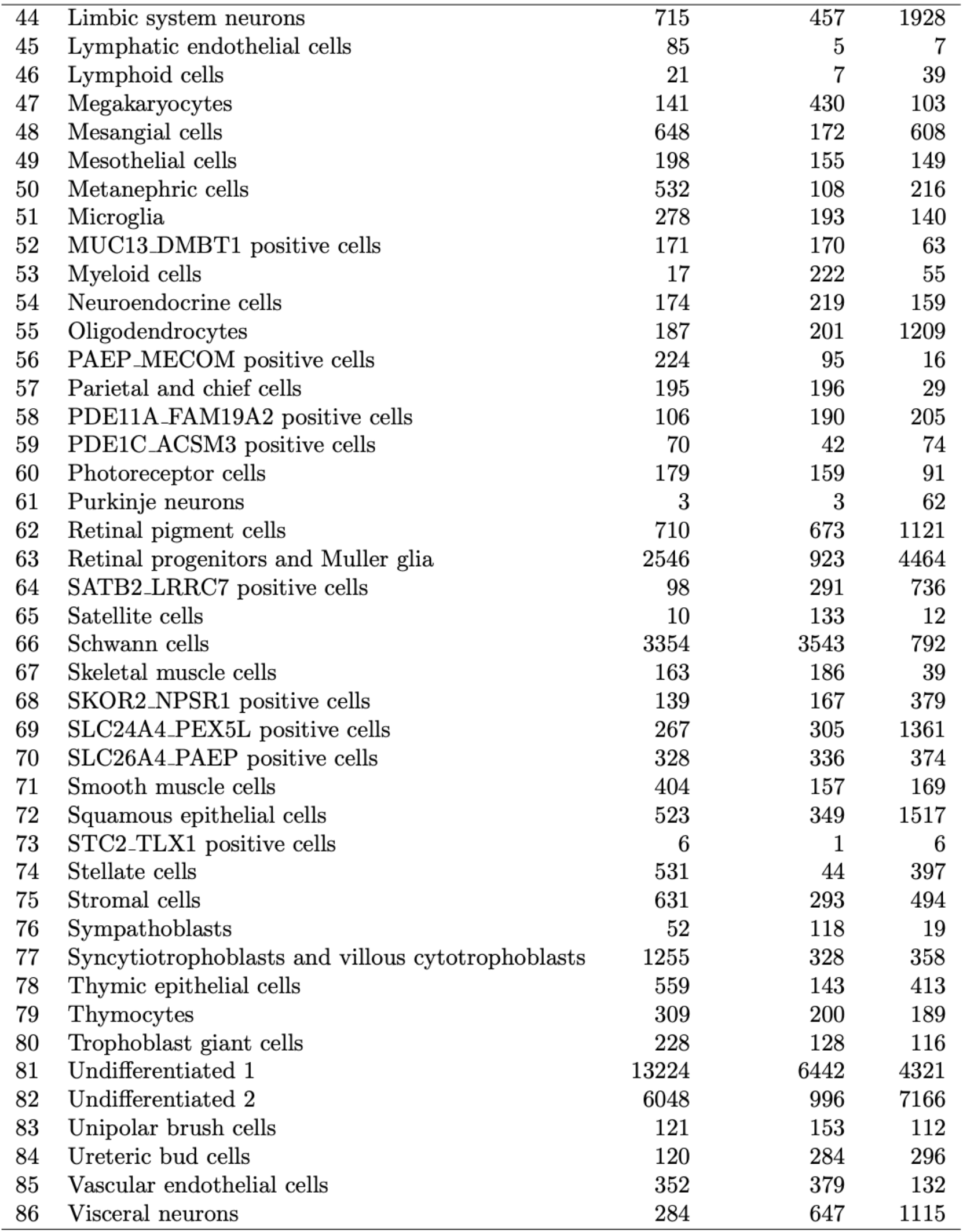
Number of cells from each species assigned to each reference cell type. Cell types are defined by the reference integration procedure described in Materials and methods.

|  | Cell type | Germ layer | Human | Chimpanzee | Hybrid |
| --- | --- | --- | --- | --- | --- |
| 1 | Acinar cells | Endoderm | 122 | 161 | 86 |
| 2 | Adrenocortical cells | Mesoderm | 163 | 145 | 59 |
| 3 | AFP_ALB positive cells | Endoderm | 465 | 18 | 384 |
| 4 | Amacrine cells | Ectoderm | 242 | 140 | 794 |
| 5 | Antigen presenting cells | Mesoderm | 57 | 32 | 19 |
| 6 | Astrocytes | Ectoderm | 1430 | 859 | 8000 |
| 7 | Bipolar cells | Ectoderm | 136 | 137 | 206 |
| 8 | Bronchiolar and alveolar epithelial cells | Endoderm | 187 | 443 | 168 |
| 9 | Cardiomyocytes | Mesoderm | 846 | 180 | 995 |
| 10 | CCL19_CCL21 positive cells | Mesoderm | 65 | 18 | 17 |
| 11 | Chromaffin cells | Ectoderm | 42 | 124 | 581 |
| 12 | Ciliated epithelial cells | Ectoderm | 319 | 723 | 407 |
| 13 | CLC_IL5RA positive cells | Mesoderm | 82 | 33 | 19 |
| 14 | Corneal and conjunctival epithelial cells | Ectoderm | 345 | 86 | 385 |
| 15 | CSH1_CSH2 positive cells | Trophoblast | 10 | 4 | 8 |
| 16 | Ductal cells | Endoderm | 558 | 129 | 239 |
| 17 | Early Ectoderm 1 | Ectoderm | 14187 | 7751 | 7865 |
| 18 | Early Ectoderm 2 | Ectoderm | 6876 | 6135 | 4491 |
| 19 | Early Ectoderm 3 | Ectoderm | 645 | 469 | 176 |
| 20 | Early Ectoderm 4 | Ectoderm | 99 | 26 | 132 |
| 21 | Early Endoderm 1 | Endoderm | 537 | 486 | 1256 |
| 22 | Early Endoderm 2 | Endoderm | 532 | 13 | 155 |
| 23 | Early Mesoderm 1 | Mesoderm | 761 | 82 | 441 |
| 24 | ELF3_AGBL2 positive cells | Mesoderm | 103 | 40 | 73 |
| 25 | Endocardial cells | Mesoderm | 123 | 11 | 19 |
| 26 | ENS glia | Ectoderm | 245 | 611 | 90 |
| 27 | ENS neurons | Ectoderm | 649 | 410 | 229 |
| 28 | Epicardial fat cells | Mesoderm | 467 | 148 | 572 |
| 29 | Erythroblasts | Mesoderm | 143 | 139 | 52 |
| 30 | Excitatory neurons | Ectoderm | 18 | 57 | 530 |
| 31 | Extravillous trophoblasts | Trophoblast | 143 | 79 | 12 |
| 32 | Ganglion cells | Ectoderm | 233 | 365 | 358 |
| 33 | Goblet cells | Endoderm | 650 | 135 | 206 |
| 34 | Granule neurons | Ectoderm | 216 | 212 | 729 |
| 35 | Hematopoietic stem cells | Mesoderm | 166 | 63 | 132 |
| 36 | Hepatoblasts | Endoderm | 111 | 548 | 159 |
| 37 | Horizontal cells | Ectoderm | 332 | 143 | 436 |
| 38 | IGFBP1_DKK1 positive cells | Mesoderm | 1123 | 274 | 1237 |
| 39 | Inhibitory interneurons | Ectoderm | 55 | 108 | 435 |
| 40 | Inhibitory neurons | Ectoderm | 22 | 68 | 1190 |
| 41 | Intestinal epithelial cells | Endoderm | 677 | 15 | 289 |
| 42 | Islet endocrine cells | Endoderm | 1087 | 1443 | 403 |
| 43 | Lens fibre cells | Ectoderm | 73 | 415 | 89 |

|  | Celltype | Human | Chimpanzee | Hybrid |
| --- | --- | --- | --- | --- |
| 1 | Acinar cells | 122 | 161 | 86 |
| 2 | Adrenocortical cells | 163 | 145 | 59 |
| 3 | AFP_ALB positive cells | 465 | 18 | 384 |
| 4 | Amacrine cells | 242 | 140 | 794 |
| 5 | Antigen presenting cells | 57 | 32 | 19 |
| 6 | Astrocytes | 1430 | 859 | 8000 |
| 7 | Bipolar cells | 136 | 137 | 206 |
| 8 | Bronchiolar and alveolar epithelial cells | 187 | 443 | 168 |
| 9 | Cardiomyocytes | 846 | 180 | 995 |
| 10 | CCL19_CCL21 positive cells | 65 | 18 | 17 |
| 11 | Chromaffin cells | 42 | 124 | 581 |
| 12 | Ciliated epithelial cells | 319 | 723 | 407 |
| 13 | CLC_IL5RA positive cells | 82 | 33 | 19 |
| 14 | Corneal and conjunctival epithelial cells | 345 | 86 | 385 |
| 15 | CSH1_CSH2 positive cells | 10 | 4 | 8 |
| 16 | Ductal cells | 558 | 129 | 239 |
| 17 | Early Ectoderm 1 | 14187 | 7751 | 7865 |
| 18 | Early Ectoderm 2 | 6876 | 6135 | 4491 |
| 19 | Early Ectoderm 3 | 645 | 469 | 176 |
| 20 | Early Ectoderm 4 | 99 | 26 | 132 |
| 21 | Early Endoderm 1 | 537 | 486 | 1256 |
| 22 | Early Endoderm 2 | 532 | 13 | 155 |
| 23 | Early Mesoderm 1 | 761 | 82 | 441 |
| 24 | ELF3_AGBL2 positive cells | 103 | 40 | 73 |
| 25 | Endocardial cells | 123 | 11 | 19 |
| 26 | ENS glia | 245 | 611 | 90 |
| 27 | ENS neurons | 649 | 410 | 229 |
| 28 | Epicardial fat cells | 467 | 148 | 572 |
| 29 | Erythroblasts | 143 | 139 | 52 |
| 30 | Excitatory neurons | 18 | 57 | 530 |
| 31 | Extravillous trophoblasts | 143 | 79 | 12 |
| 32 | Ganglion cells | 233 | 365 | 358 |
| 33 | Goblet cells | 650 | 135 | 206 |
| 34 | Granule neurons | 216 | 212 | 729 |
| 35 | Hematopoietic stem cells | 166 | 63 | 132 |
| 36 | Hepatoblasts | 111 | 548 | 159 |
| 37 | Horizontal cells | 332 | 143 | 436 |
| 38 | IGFBP1_DKK1 positive cells | 1123 | 274 | 1237 |
| 39 | Inhibitory interneurons | 55 | 108 | 435 |
| 40 | Inhibitory neurons | 22 | 68 | 1190 |
| 41 | Intestinal epithelial cells | 677 | 15 | 289 |
| 42 | Islet endocrine cells | 1087 | 1443 | 403 |
| 43 | Lens fibre cells | 73 | 415 | 89 |
| 44 | Limbic system neurons | Ectoderm | 715 | 457 | 1928 |
| 45 | Lymphatic endothelial cells | Mesoderm | 85 | 5 | 7 |
| 46 | Lymphoid cells | Mesoderm | 21 | 7 | 39 |
| 47 | Megakaryocytes | Mesoderm | 141 | 430 | 103 |
| 48 | Mesangial cells | Mesoderm | 648 | 172 | 608 |
| 49 | Mesothelial cells | Mesoderm | 198 | 155 | 149 |
| 50 | Metanephric cells | Mesoderm | 532 | 108 | 216 |
| 51 | Microglia | Mesoderm | 278 | 193 | 140 |
| 52 | MUC13_DMBT1 positive cells | Endoderm | 171 | 170 | 63 |
| 53 | Myeloid cells | Mesoderm | 17 | 222 | 55 |
| 54 | Neuroendocrine cells | Ectoderm | 174 | 219 | 159 |
| 55 | Oligodendrocytes | Ectoderm | 187 | 201 | 1209 |
| 56 | PAEP_MECOM positive cells | Mesoderm | 224 | 95 | 16 |
| 57 | Parietal and chief cells | Endoderm | 195 | 196 | 29 |
| 58 | PDE11A_FAM19A2 positive cells | Ectoderm | 106 | 190 | 205 |
| 59 | PDE1C_ACSM3 positive cells | Ectoderm | 70 | 42 | 74 |
| 60 | Photoreceptor cells | Ectoderm | 179 | 159 | 91 |
| 61 | Purkinje neurons | Ectoderm | 3 | 3 | 62 |
| 62 | Retinal pigment cells | Ectoderm | 710 | 673 | 1121 |
| 63 | Retinal progenitors and Muller glia | Ectoderm | 2546 | 923 | 4464 |
| 64 | SATB2_LRRC7 positive cells | Ectoderm | 98 | 291 | 736 |
| 65 | Satellite cells | Mesoderm | 10 | 133 | 12 |
| 66 | Schwann cells | Ectoderm | 3354 | 3543 | 792 |
| 67 | Skeletal muscle cells | Mesoderm | 163 | 186 | 39 |
| 68 | SKOR2_NPSR1 positive cells | Ectoderm | 139 | 167 | 379 |
| 69 | SLC24A4_PEX5L positive cells | Ectoderm | 267 | 305 | 1361 |
| 70 | SLC26A4_PAEP positive cells | Ectoderm | 328 | 336 | 374 |
| 71 | Smooth muscle cells | Mesoderm | 404 | 157 | 169 |
| 72 | Squamous epithelial cells | Ectoderm | 523 | 349 | 1517 |
| 73 | STC2_TLX1 positive cells | Mesoderm | 6 | 1 | 6 |
| 74 | Stellate cells | Mesoderm | 531 | 44 | 397 |
| 75 | Stromal cells | Mesoderm | 631 | 293 | 494 |
| 76 | Sympathoblasts | Ectoderm | 52 | 118 | 19 |
| 77 | Syncytiotrophoblasts and villous cytotrophoblasts | Trophoblast | 1255 | 328 | 358 |
| 78 | Thymic epithelial cells | Endoderm | 559 | 143 | 413 |
| 79 | Thymocytes | Mesoderm | 309 | 200 | 189 |
| 80 | Trophoblast giant cells | Trophoblast | 228 | 128 | 116 |
| 81 | Undifferentiated 1 | Undifferentiated | 13224 | 6442 | 4321 |
| 82 | Undifferentiated 2 | Undifferentiated | 6048 | 996 | 7166 |
| 83 | Unipolar brush cells | Ectoderm | 121 | 153 | 112 |
| 84 | Ureteric bud cells | Ectoderm | 120 | 284 | 296 |
| 85 | Vascular endothelial cells | Mesoderm | 352 | 379 | 132 |
| 86 | Visceral neurons | Ectoderm | 284 | 647 | 1115 |

|  | Celltype | Human | Chimpanzee | Hybrid |
| --- | --- | --- | --- | --- |
| 44 | Limbic system neurons | 715 | 457 | 1928 |
| 45 | Lymphatic endothelial cells | 85 | 5 | 7 |
| 46 | Lymphoid cells | 21 | 7 | 39 |
| 47 | Megakaryocytes | 141 | 430 | 103 |
| 48 | Mesangial cells | 648 | 172 | 608 |
| 49 | Mesothelial cells | 198 | 155 | 149 |
| 50 | Metanephric cells | 532 | 108 | 216 |
| 51 | Microglia | 278 | 193 | 140 |
| 52 | MUC13.DMBT1 positive cells | 171 | 170 | 63 |
| 53 | Myeloid cells | 17 | 222 | 55 |
| 54 | Neuroendocrine cells | 174 | 219 | 159 |
| 55 | Oligodendrocytes | 187 | 201 | 1209 |
| 56 | PAEP.MECOM positive cells | 224 | 95 | 16 |
| 57 | Parietal and chief cells | 195 | 196 | 29 |
| 58 | PDE11A.FAM19A2 positive cells | 106 | 190 | 205 |
| 59 | PDE1C.ACSM3 positive cells | 70 | 42 | 74 |
| 60 | Photoreceptor cells | 179 | 159 | 91 |
| 61 | Purkinje neurons | 3 | 3 | 62 |
| 62 | Retinal pigment cells | 710 | 673 | 1121 |
| 63 | Retinal progenitors and Muller glia | 2546 | 923 | 4464 |
| 64 | SATB2.LRRC7 positive cells | 98 | 291 | 736 |
| 65 | Satellite cells | 10 | 133 | 12 |
| 66 | Schwann cells | 3354 | 3543 | 792 |
| 67 | Skeletal muscle cells | 163 | 186 | 39 |
| 68 | SKOR2.NPSR1 positive cells | 139 | 167 | 379 |
| 69 | SLC24A4.PEX5L positive cells | 267 | 305 | 1361 |
| 70 | SLC26A4.PAEP positive cells | 328 | 336 | 374 |
| 71 | Smooth muscle cells | 404 | 157 | 169 |
| 72 | Squamous epithelial cells | 523 | 349 | 1517 |
| 73 | STC2.TLX1 positive cells | 6 | 1 | 6 |
| 74 | Stellate cells | 531 | 44 | 397 |
| 75 | Stromal cells | 631 | 293 | 494 |
| 76 | Sympathoblasts | 52 | 118 | 19 |
| 77 | Syncytiotrophoblasts and villous cytotrophoblasts | 1255 | 328 | 358 |
| 78 | Thymic epithelial cells | 559 | 143 | 413 |
| 79 | Thymocytes | 309 | 200 | 189 |
| 80 | Trophoblast giant cells | 228 | 128 | 116 |
| 81 | Undifferentiated 1 | 13224 | 6442 | 4321 |
| 82 | Undifferentiated 2 | 6048 | 996 | 7166 |
| 83 | Unipolar brush cells | 121 | 153 | 112 |
| 84 | Ureteric bud cells | 120 | 284 | 296 |
| 85 | Vascular endothelial cells | 352 | 379 | 132 |
| 86 | Visceral neurons | 284 | 647 | 1115 |

**Table S4.** Number of differentially expressed genes by cell type. The number of differentially expressed cells calculated with DREAM at 5% false discovery rate, Cormotif and 95% posterior probability of DE, or in tetraploid hybrid cells using the Wilcoxon signed rank test at 5% false discovery rate.

|  | Cell Type | DREAM | CorMotif | Hybrid |
| --- | --- | --- | --- | --- |
| 1 | Early Ectoderm 2 | 2034 | 6193 | 2034 |
| 2 | Hepatoblasts | 1200 | 3911 | 1200 |
| 3 | Cardiomyocytes | 563 | 2062 | 563 |
| 4 | Syncytiotrophoblasts and villous cytotrophoblasts | 3019 | 6359 | 3019 |
| 5 | Schwann cells | 1449 | 3821 | 1449 |
| 6 | Undifferentiated 1 | 1682 | 7095 | 1682 |
| 7 | Ciliated epithelial cells | 794 | 3224 | 794 |
| 8 | IGFBP1_DKK1 positive cells | 278 | 1526 | 278 |
| 9 | Undifferentiated 2 | 2073 | 5320 | 2073 |
| 10 | Bipolar cells | 511 | 2768 | 511 |
| 11 | Metanephric cells | 105 | 801 | 105 |
| 12 | Trophoblast giant cells | 788 | 4134 | 788 |
| 13 | SLC24A4_PEX5L positive cells | 350 | 1566 | 350 |
| 14 | Early Ectoderm 1 | 3712 | 6709 | 3712 |
| 15 | Squamous epithelial cells | 476 | 3088 | 476 |
| 16 | Retinal progenitors and Muller glia | 3135 | 5794 | 3135 |
| 17 | Thymocytes | 1282 | 5115 | 1282 |
| 18 | Astrocytes | 3681 | 6640 | 3681 |
| 19 | Adrenocortical cells | 346 | 4188 | 346 |
| 20 | Stromal cells | 799 | 3424 | 799 |
| 21 | Parietal and chief cells | 540 | 4398 | 540 |
| 22 | SKOR2_NPSR1 positive cells | 463 | 2896 | 463 |
| 23 | Goblet cells | 1532 | 3139 | 1532 |
| 24 | Mesangial cells | 114 | 1188 | 114 |
| 25 | Epicardial fat cells | 515 | 2232 | 515 |
| 26 | PDE11A_FAM19A2 positive cells | 525 | 2536 | 525 |
| 27 | ENS neurons | 614 | 2280 | 614 |
| 28 | Smooth muscle cells | 367 | 2484 | 367 |
| 29 | Early Ectoderm 3 | 492 | 2470 | 492 |
| 30 | Vascular endothelial cells | 993 | 3410 | 993 |
| 31 | Limbic system neurons | 753 | 2050 | 753 |
| 32 | Early Endoderm 1 | 433 | 4791 | 433 |
| 33 | ENS glia | 516 | 2251 | 516 |
| 34 | Oligodendrocytes | 493 | 2866 | 493 |
| 35 | Hematopoietic stem cells | 105 | 911 | 105 |
| 36 | Ureteric bud cells | 488 | 2667 | 488 |
| 37 | Visceral neurons | 160 | 1726 | 160 |
| 38 | PAEP_MECOM positive cells | 569 | 3695 | 569 |
| 39 | Retinal pigment cells | 1451 | 4251 | 1451 |
| 40 | Early Ectoderm 4 | 0 | 413 | 0 |
| 41 | Amacrine cells | 138 | 1104 | 138 |
| 42 | Ductal cells | 888 | 1976 | 888 |
| 43 | Early Mesoderm 1 | 89 | 625 | 89 |
| 44 | MUC13.DMBT1 positive cells | 1358 | 3324 | 1358 |
| 45 | Bronchiolar and alveolar epithelial cells | 879 | 3950 | 879 |
| 46 | Ganglion cells | 381 | 1874 | 381 |
| 47 | Thymic epithelial cells | 353 | 1860 | 353 |
| 48 | Horizontal cells | 197 | 1506 | 197 |
| 49 | Islet endocrine cells | 2218 | 4293 | 2218 |
| 50 | Granule neurons | 450 | 1618 | 450 |
| 51 | Unipolar brush cells | 292 | 1103 | 292 |
| 52 | Corneal and conjunctival epithelial cells | 91 | 604 | 91 |
| 53 | Mesothelial cells | 1452 | 3120 | 1452 |
| 54 | Antigen presenting cells | 42 | 779 | 42 |
| 55 | Neuroendocrine cells | 340 | 1760 | 340 |
| 56 | Stellate cells | 323 | 1490 | 323 |
| 57 | PDE1C_ACSM3 positive cells | 206 | 1679 | 206 |
| 58 | Photoreceptor cells | 583 | 2596 | 583 |
| 59 | Skeletal muscle cells | 436 | 4718 | 436 |
| 60 | Microglia | 2233 | 5124 | 2233 |
| 61 | SLC26A4.PAEP positive cells | 1638 | 4633 | 1638 |
| 62 | Erythroblasts | 656 | 4622 | 656 |
| 63 | Megakaryocytes | 2118 | 5403 | 2118 |
| 64 | Chromaffin cells | 2 | 27 | 2 |
| 65 | Extravillous trophoblasts | 609 | 2950 | 609 |
| 66 | ELF3_AGBL2 positive cells | 7 | 410 | 7 |
| 67 | CLC_IL5RA positive cells | 0 | 429 | 0 |
| 68 | Inhibitory interneurons | 33 | 349 | 33 |
| 69 | Lens fibre cells | 463 | 2558 | 463 |
| 70 | Sympathoblasts | 522 | 4216 | 522 |
| 71 | SATB2_LRRC7 positive cells | 34 | 1119 | 34 |
| 72 | Acinar cells | 1509 | 4764 | 1509 |

**Table S5.** Gene Ontology enrichment in genes not differentially expressed in any cell type. The gene set was filtered for genes that had at least 100 UMIs detected across all individuals and replicates in at least 10 cell types. The top 20 terms are provided.

|  | GO.ID | Term | p-value |
| --- | --- | --- | --- |
| 1 | GO:0008380 | RNA splicing | 0.00042 |
| 2 | GO:0045931 | positive regulation of mitotic cell cycl... | 0.00044 |
| 3 | GO:0006397 | mRNA processing | 0.00049 |
| 4 | GO:0032007 | negative regulation of TOR signaling | 0.00197 |
| 5 | GO:0006513 | protein monoubiquitination | 0.00212 |
| 6 | GO:0043966 | histone H3 acetylation | 0.00294 |
| 7 | GO:0051098 | regulation of binding | 0.00318 |
| 8 | GO:0016236 | macroautophagy | 0.00392 |
| 9 | GO:0016032 | viral process | 0.00433 |
| 10 | GO:0000278 | mitotic cell cycle | 0.00515 |
| 11 | GO:0016925 | protein sumoylation | 0.00552 |
| 12 | GO:0032663 | regulation of interleukin-2 production | 0.00832 |
| 13 | GO:0051028 | mRNA transport | 0.00951 |
| 14 | GO:0006367 | transcription initiation from RNA polyme... | 0.01289 |
| 15 | GO:0043161 | proteasome-mediated ubiquitin-dependent ... | 0.01318 |
| 16 | GO:0000338 | protein deneddylation | 0.01594 |
| 17 | GO:2000757 | negative regulation of peptidyl-lysine a... | 0.01594 |
| 18 | GO:0043967 | histone H4 acetylation | 0.01598 |
| 19 | GO:0006606 | protein import into nucleus | 0.01610 |
| 20 | GO:0016567 | protein ubiquitination | 0.01703 |

**Table S6.** Gene Ontology enrichment for *trans* genes. We ranked genes according to the mean *trans* proportion across all cell types with DE and ran gene set enrichment against Gene Ontology Biological Process terms. The top 20 terms are provided.

|  | pathway | p-value | Normalized enrichment score |
| --- | --- | --- | --- |
| 1 | TUBE_MORPHOGENESIS | 9.8E-10 | 1.8 |
| 2 | TUBE_DEVELOPMENT | 2.2E-09 | 1.8 |
| 3 | REGULATION_OF_MULTICELLULAR_ORGANISMAL_DEVELOPMENT | 9.8E-09 | 1.7 |
| 4 | CELL_FATE_COMMITMENT | 1.1E-08 | 2.3 |
| 5 | TISSUE_DEVELOPMENT | 2.9E-08 | 1.6 |
| 6 | ANIMAL_ORGAN_MORPHOGENESIS | 4.2E-08 | 1.8 |
| 7 | ANATOMICAL_STRUCTURE_FORMATION_INVOLVED_IN_MORP... | 1.3E-07 | 1.7 |
| 8 | EMBRYO_DEVELOPMENT | 2.2E-07 | 1.7 |
| 9 | ENZYME_LINKED_RECEPTOR_PROTEIN_SIGNALING_PATHWAY | 2.2E-07 | 1.7 |
| 10 | BIOLOGICAL_ADHESION | 2.4E-07 | 1.6 |
| 11 | REGULATION_OF_CELL_POPULATION_PROLIFERATION | 6.0E-07 | 1.6 |
| 12 | EMBRYONIC_ORGAN_DEVELOPMENT | 6.5E-07 | 1.9 |
| 13 | CELL_CYCLE | 1.4E-06 | 1.5 |
| 14 | EMBRYONIC_MORPHOGENESIS | 2.0E-06 | 1.8 |
| 15 | CIRCULATORY_SYSTEM_DEVELOPMENT | 2.5E-06 | 1.6 |
| 16 | EPITHELIUM_DEVELOPMENT | 6.1E-06 | 1.6 |
| 17 | REGULATION_OF_CELL_DIFFERENTIATION | 6.1E-06 | 1.5 |
| 18 | PATTERN_SPECIFICATION_PROCESS | 1.2E-05 | 1.9 |
| 19 | POSITIVE_REGULATION_OF_CELL_POPULATION_PROLIFER... | 1.3E-05 | 1.7 |
| 20 | POSITIVE_REGULATION_OF_DEVELOPMENTAL_PROCESS | 1.5E-05 | 1.5 |

**Table S7.**
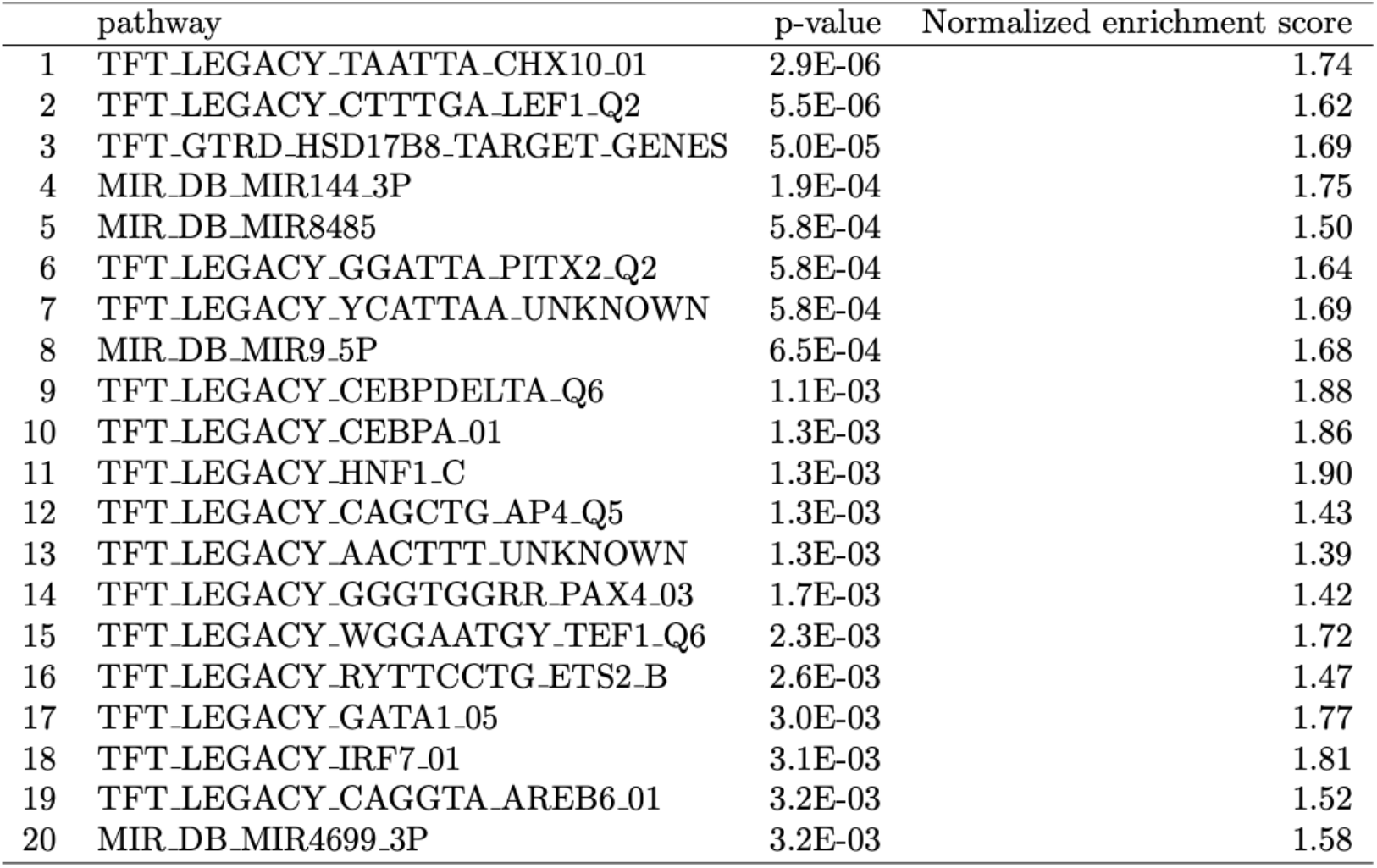
Transcription factor target enrichment for *trans* genes. We ranked genes according to the mean *trans* proportion across all cell types with DE and ran gene set enrichment against transcription factor targets. The top 20 terms are provided.

### Data S1-S2

**Data S1. Results of DREAM, Cormotif, Hybrid DE, and cis proportion**. This supplementary data file is a flat tab-separated text file with the following columns defined:

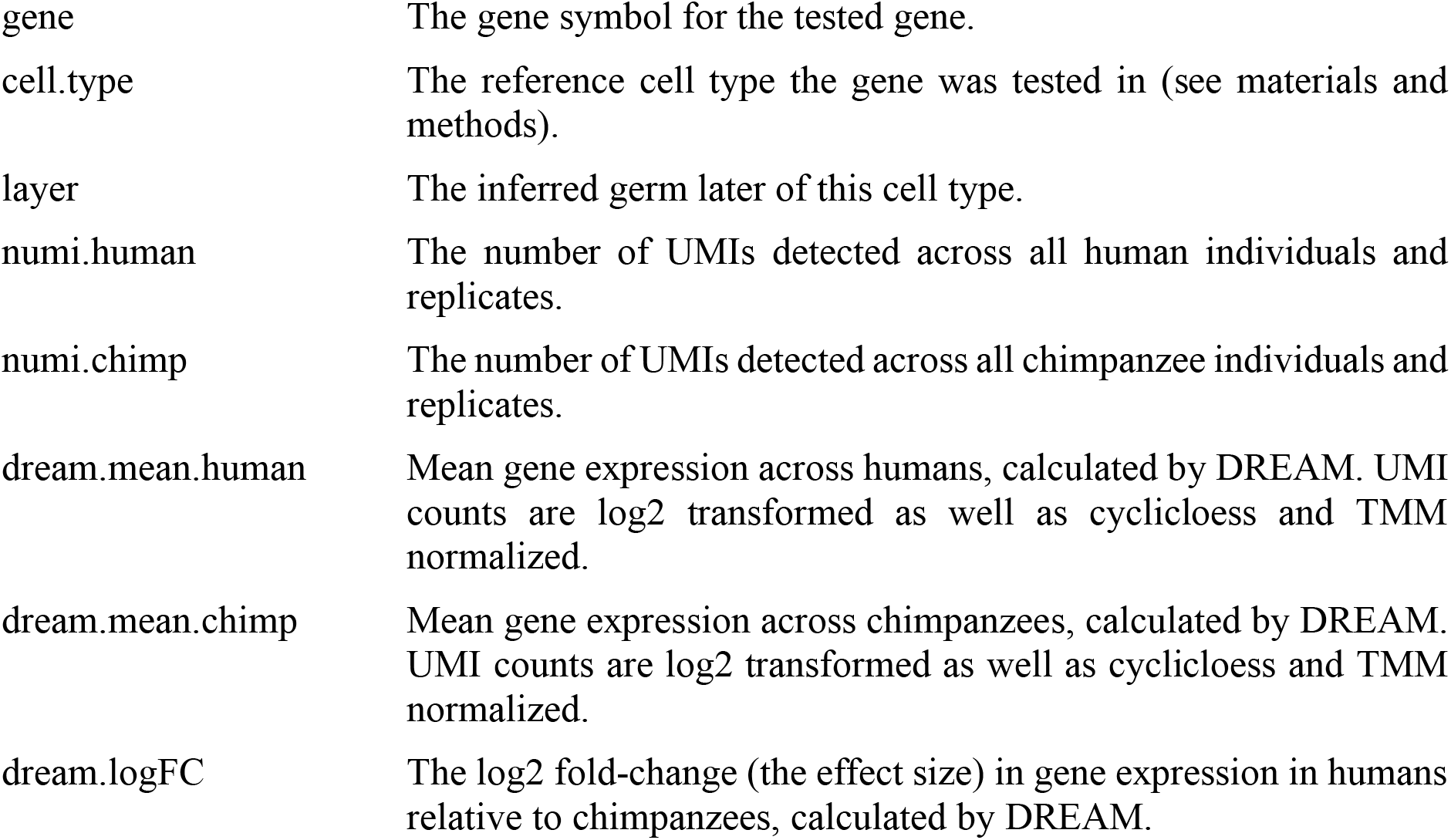

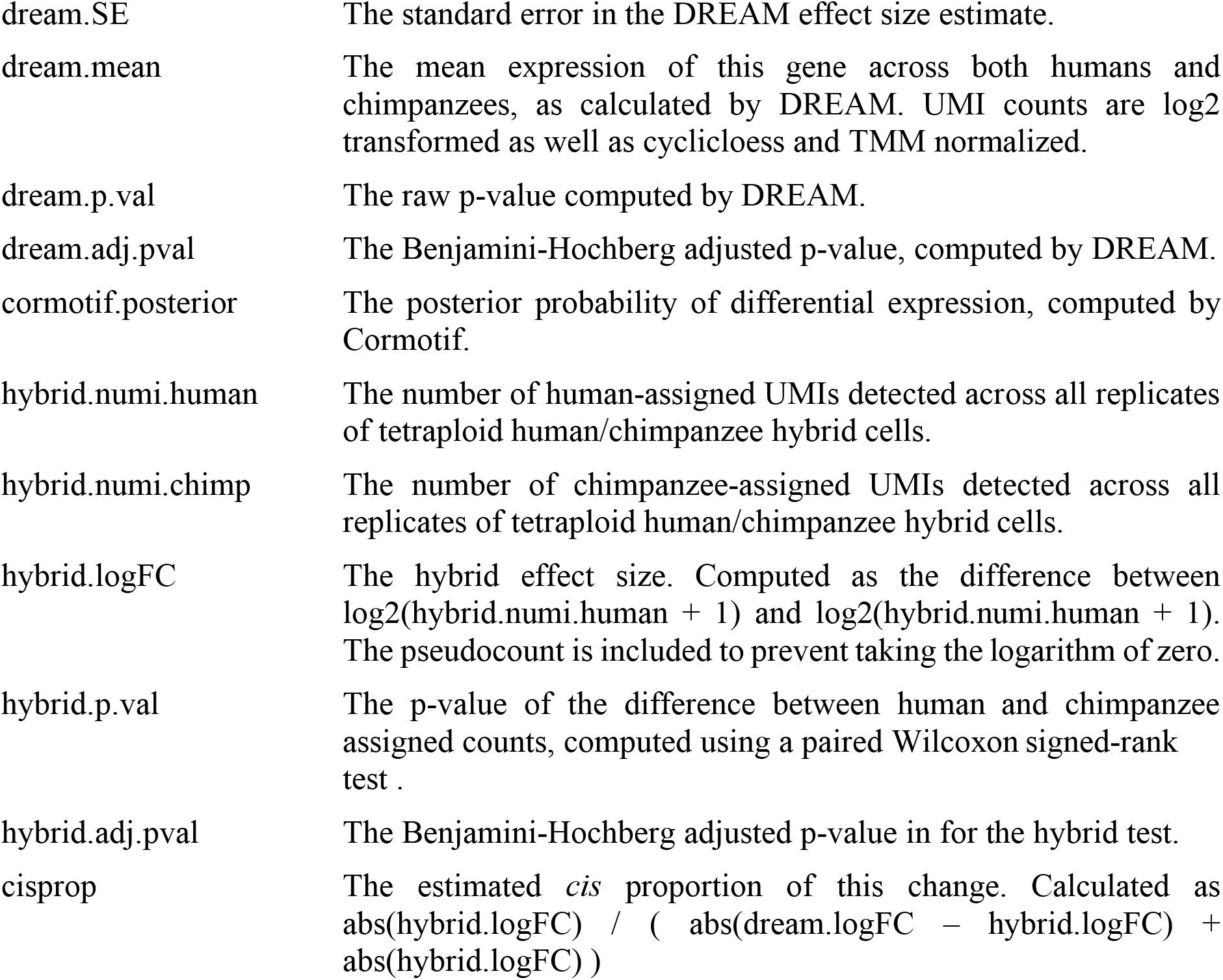

**Data S2. Genes with cell-type restricted differential expression**. Raw text file containing the gene symbol of each gene with cell-type-restricted differential expression. One gene is listed per line.

